# Targeting protein-ligand neosurfaces using a generalizable deep learning approach

**DOI:** 10.1101/2024.03.25.585721

**Authors:** Anthony Marchand, Stephen Buckley, Arne Schneuing, Martin Pacesa, Pablo Gainza, Evgenia Elizarova, Rebecca M. Neeser, Pao-Wan Lee, Luc Reymond, Maddalena Elia, Leo Scheller, Sandrine Georgeon, Joseph Schmidt, Philippe Schwaller, Sebastian J. Maerkl, Michael Bronstein, Bruno E. Correia

**Author notes:** These authors contributed equally to this work (Anthony Marchand, Stephen Buckley, Arne Schneuing).

## Abstract

Molecular recognition events between proteins drive biological processes in living systems. However, higher levels of mechanistic regulation have emerged, where protein-protein interactions are conditioned to small molecules. Here, we present a computational strategy for the design of proteins that target neosurfaces, i.e. surfaces arising from protein-ligand complexes. To do so, we leveraged a deep learning approach based on learned molecular surface representations and experimentally validated binders against three drug-bound protein complexes. Remarkably, surface fingerprints trained only on proteins can be applied to neosurfaces emerging from small molecules, serving as a powerful demonstration of generalizability that is uncommon in deep learning approaches. The designed chemically-induced protein interactions hold the potential to expand the sensing repertoire and the assembly of new synthetic pathways in engineered cells.

## Main

Protein-protein interactions (PPIs) play an essential role in healthy cell homeostasis, but are also involved in numerous diseases(*1*, *2*). For this reason, several therapies targeting PPIs have been developed over the last decades and multiple computational tools have been recently proposed to design novel protein interactions(*3*). The governing principles determining the propensity of proteins to form interactions are intricate due to the interplay of several contributions, such as geometric and chemical complementarity, dynamics, and solvent interactions. Therefore it remains challenging to predict and design novel PPIs, especially in the absence of evolutionary constraints. Native PPIs can also be controlled by additional regulatory layers such as allostery(*4*), post-translational modifications(*5*), or direct ligand binding(*6*, *7*). Compound-bound surfaces, which we refer to as neosurfaces, are one of the most fascinating and challenging types of molecular recognition instances, where relatively minor changes at the protein binding site can have a large impact on binding affinities. The interest in such interactions has been fueled by the development of new drug modalities; specifically molecular glues that form neosurfaces to trigger protein interactions for degradation and other applications(*8*, *9*), thus representing a promising route for the development of innovative therapeutics.

In synthetic biology, molecular components that rely on small molecule-induced neosurfaces have been used to engineer chemically-responsive systems with precise spatio-temporal control of cellular activities(*10*). Small molecule triggers have been used to both induce and disrupt PPIs, thereby functioning as ON or OFF switches for engineered cellular functions(*10*– *12*). There are several practical advantages in using small molecules as triggers due to their simple administration, biodistribution, cell permeability, safety, and high affinity and specificity to their target proteins. Protein-based switches controlled by small molecules have already been applied to regulate transcription(*13*, *14*), protein degradation(*15–17*), and protein localization(*18–20*), among many other applications. In addition to their use in basic research, engineering molecular switches is becoming a more common mechanism of controlling protein-based and cellular therapeutics, whose activity may have to be regulated to mitigate potentially dangerous side effects(*11*, *21*, *22*). While several chemically-disruptable heterodimer (OFF-switch) systems have been proposed(*10*, *11*, *21*), computationally designed chemically-induced dimerization (CID, ON-switch) systems remain challenging due to the complexity of modeling neosurfaces. Previous attempts at designing CID systems primarily relied on experimental methods(*10*, *13*, *23*, *24*) and, despite the emergence of artificial intelligence and numerous computational tools, only few tools can generalize to both proteins and small molecules as a target for protein design, resulting in a lack of suitable approaches for the design of novel chemically-induced PPIs. Computational methods to design novel CIDs mostly relied on transplanting an existing drug binding site to a known heterodimer interface(*25*) or using docking of putative pre-existing proteins (i.e. scaffolds) followed by interface optimization(*26*). However, these approaches can face limitations such as the risk of drug-independent dimerization, the lack of suitable scaffold proteins for design, or the extensive need for *in vitro* maturation techniques.

We recently reported a geometric deep learning-based framework called MaSIF (Molecular Surface Interaction Fingerprinting)(*27*) for the study of protein surface features, and for the design of novel protein-protein interactions(*28*). In this study, we aim to test whether our surface-centric approach can generalize to non-protein ligands without additional training data by using a higher-level representation, namely the geometric and chemical features found on the molecular surface. To do so, we designed site-specific binders that target neosurfaces composed of a small molecule ligand and protein surface moieties, resulting in *de novo* ligand-dependent protein interactions. We successfully designed and characterized novel protein binders recognizing the B-cell lymphoma 2 (Bcl2) protein in complex with the clinically-approved inhibitor Venetoclax(*29*), the progesterone-binding antibody DB3 in complex with its ligand(*30*), and finally the peptide deformylase 1 (PDF1) protein from *Pseudomonas aeruginosa* in complex with the antibiotic Actinonin(*31*). Lastly, we show that such ligand-controlled systems can be utilized in both *in vitro* and cellular contexts for a range of synthetic biology applications, unlocking possibilities for the development and regulation of novel therapeutic approaches.

### MaSIF captures interaction propensities of neosurfaces

Within our geometric deep learning framework, MaSIF(*27*), we previously developed two applications: i) *MaSIF-site* to accurately predict regions of the protein surface with a high propensity to form an interface with another protein and, ii) *MaSIF-search* to rapidly find and dock protein partners based on complementary surface patches. In *MaSIF-search*, we extract surface patch descriptors (“fingerprints”), so that patches with complementary geometry and chemistry have similar fingerprints, whereas non-interacting patches have low fingerprint similarity. Surface fingerprints allow to perform an initial ultra-fast search in an alignment-free manner using the Euclidean distances between them. Patches with fingerprint distances below a threshold are then further aligned in 3D and scored with an interface-post alignment (IPA) score to refine the selection.

In its initial conception, MaSIF only considered canonical amino acids as part of the protein molecular surface and was not compatible with small molecules, glycans, and other ligands. Thus, we present here *MaSIF-neosurf* to incorporate small molecules as part of the molecular surface representation of the target protein to predict interfaces and partners based on the neosurface fingerprints (Fig. 1A, see Methods). MaSIF was initially trained to operate on general chemical and geometric surface properties of biomolecules, while abstracting the underlying structure. Thus, it is not restricted to only protein surfaces, but should in principle also capture the surface patterns arising from other non-protein surfaces. Upon generation of the molecular surface of the protein-drug complex, *MaSIF-neosurf* computes the two geometric features: shape index(*32*) and distance-dependent curvature(*33*). In addition, three chemical features are also used: Poisson-Boltzmann electrostatics, which can be computed directly from the small molecule, and hydrogen bond donor/acceptor propensity(*34*) and hydrophobicity(*35–37*), for which we developed new featurizers tailored to capture the chemical properties of the small molecules (see Methods and fig. S1).

**Fig. 1:**
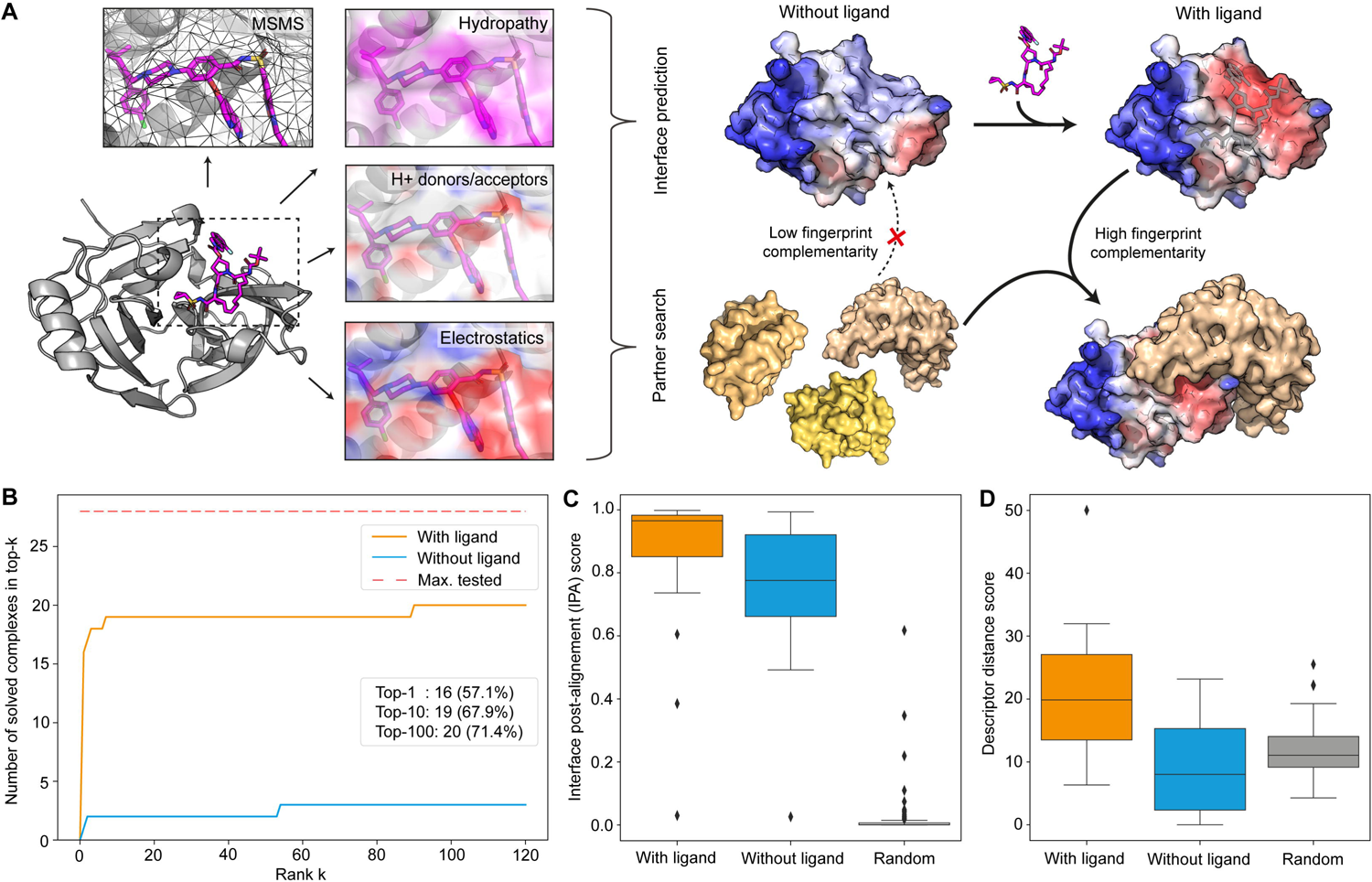
Neosurface properties are captured to identify interface sites and binding partners. **A.** Geometric and chemical features of the ligand-protein complexes are computed, including the molecular surface representation (MSMS), hydropathy score, proton donors/acceptors and Poisson-Boltzmann electrostatics. Surface features are vectorized in a descriptor (also referred to as “fingerprint”) and used by *MaSIF-neosurf* for interface propensity prediction or protein partner search. The ligand-containing fingerprint is then used to find complementary fingerprints in a patch database. **B.** Ranking predictions using *MaSIF-neosurf* on a benchmark dataset of known ternary complexes and a set of 200 decoys. Complementary partner search was performed in the presence (orange) and absence (blue) of the respective small molecule ligand. **C-D.** Interface post-alignment score (IPA; C) and descriptor distance score (see Methods; D) of the interacting complexes in the presence (orange) and absence (blue) of the drug compared to a set of random patch alignments (gray)

To assess the capabilities of *MaSIF-neosurf*, we benchmarked its performance on several ternary complexes whose interface is composed of protein and ligand surfaces. We aimed to recover known binding partners for proteins with small molecules at the binding interface. After assembling a list of 14 ligand-induced protein complexes, we split each of them into two subunits, resulting in 28 independent benchmarking cases, and processed them with and without the small molecule bound. The ligand-free protein surfaces, together with 200 decoy proteins, constitute our database, which we query with surface patches from all 28 protein-ligand complexes. Since each of the 228 protein candidates is decomposed into almost 4000 patches on average, the database represents a large search space with more than 900’000 potential binding sites. We then evaluated whether the correct binding partner is retrieved and docked in the correct rigid-body orientation. When considering the protein-ligand complex as a docking partner, *MaSIF-neosurf* recovers more than 70% of the correct binding partners and their binding poses (Fig. 1B). Only a small subset of test cases could be recovered in the absence of the ligand and the general trend is that in such cases the protein surface is a large contributor towards the overall protein interaction (fig. S2). The ability to capture the neosurface properties is further supported by an increased descriptor distance score between interacting partners (i.e. an increased complementarity between interacting fingerprints, see Methods) and an increased interface post-alignment (IPA, see Methods) score in the presence of the small molecule compared to the case without (Fig. 1C-D). Overall, *MaSIF-neosurf* captures, in many instances, features that are determinant for ligand-mediated protein interactions and, to further test its capabilities, we sought to *de novo* design this type of interactions.

### Designing novel ligand-induced protein interactions

Recently, we proposed the *MaSIF-seed* pipeline for the design of *de novo* site-specific protein binders(*28*). Given the performance of *MaSIF-seed* against multiple therapeutically relevant targets, we sought to test whether such an approach could generalize to design site-specific binders to neosurfaces composed of ligand and protein atoms. By doing so, we tackle the challenge of designing chemically controlled protein interactions and test our understanding of molecular recognition events mediated by neosurfaces. We therefore adapted our *MaSIF-seed* pipeline to our newly proposed *MaSIF-neosurf* framework (Fig. 2A). Once neosurfaces are computed for a given protein-ligand complex, we first take advantage of *MaSIF-site* to identify the regions most likely to become buried in an interface. Then an extensive fingerprint search identifies complementary structural motifs (i.e. binding seeds) from a database of ∼640’000 structural fragments (402 million surface patches/fingerprints). Therefore, by focusing on the predicted buried regions of the interface and searching for highly complementary motifs, the vast space of patches and binding motifs is quickly reduced to the most promising candidates. Finally, the top seeds are refined by sequence optimization and grafted with Rosetta(*38*) on recipient proteins (i.e. scaffolds) to stabilize the binding motif. Lastly, a final round of sequence design is performed to improve atomic contacts at the interface.

**Fig. 2:**
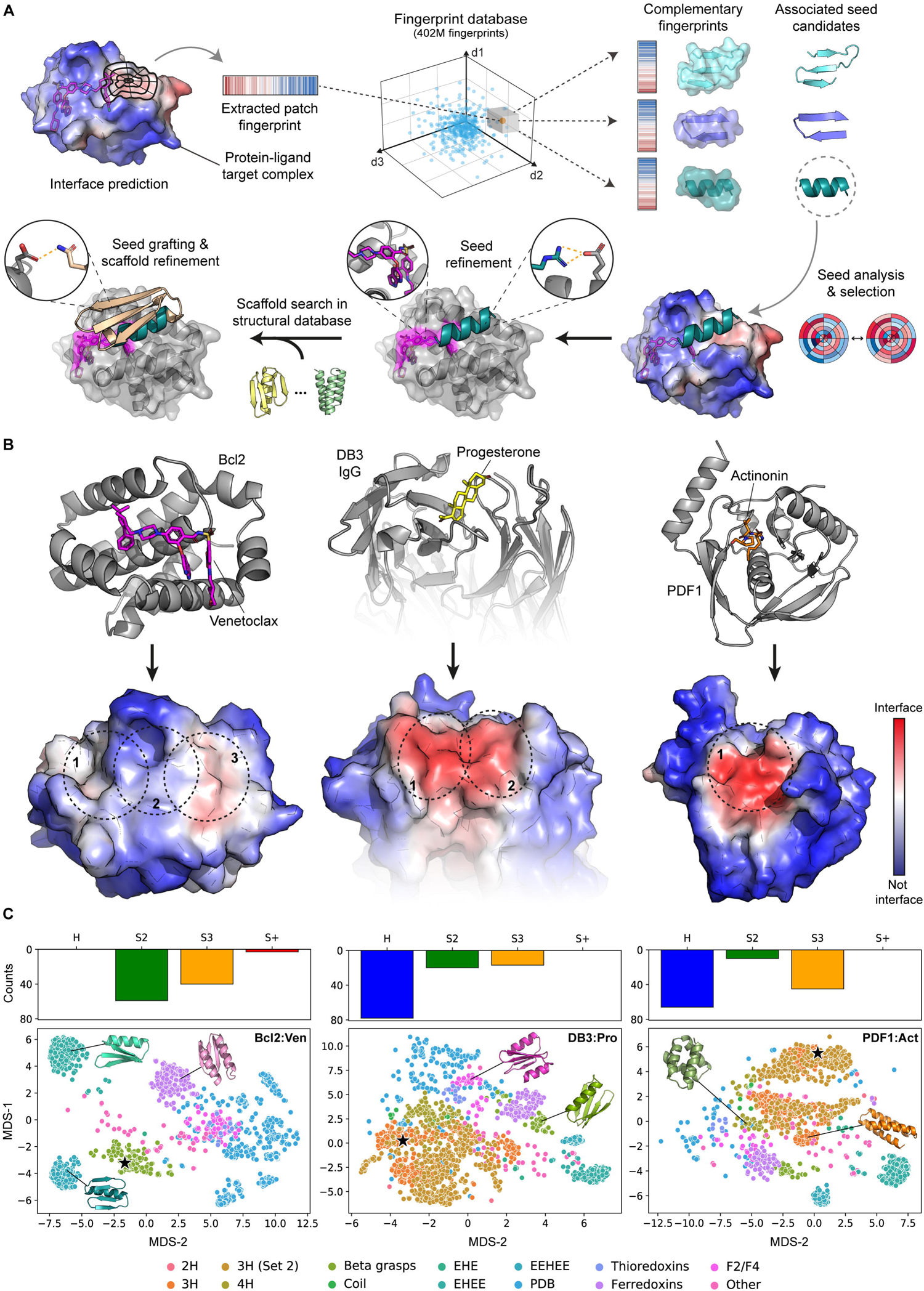
Design of ligand-induced protein interactions with *MaSIF-neosurf*. **A.** To design novel ligand-induced protein interactions, potential interface sites are first identified on the target protein-ligand complex. The corresponding patches are then used to find complementary fingerprints in a patch database. The top patches are aligned and scored to refine the selection. Associated binding motifs (seeds) undergo sequence optimization with an emphasis on designing new hydrogen bond networks with the target protein and small molecule. Seeds are then grafted on suitable scaffolds from a structural database, and the rest of the scaffold interface is redesigned using Rosetta. Finally, the top ∼2000 designs, according to different structural metrics, are selected and screened experimentally. **B.** Target candidates in complex with their respective small molecules (top row). Neosurfaces displaying their protein binding propensity (bottom row). Sites selected for binder design are highlighted with dashed circles. **C.** Seed structural diversity (top row) includes motifs that are: helical (H); two-strand beta sheets (S2); three-strand beta sheets (S3); and more complex beta sheet motifs (S+). Diversity of the ∼2000 computational designs (bottom row) mapped using multidimensional scaling (MDS) of pairwise RMSDs between all designs. Experimentally confirmed binders are highlighted with a star.

We designed ligand-dependent protein binders targeting ligand-bound proteins from different families: Bcl2 in complex with the clinically approved drug Venetoclax; an anti-progesterone antibody (DB3) in complex with its ligand; and peptide deformylase 1 (PDF1) from *P. aeruginosa* in complex with the antibiotic Actinonin (Fig 2B). We first identified a moderate to high interface propensity of these neosurfaces with *MaSIF-neosurf*, selected 1 to 3 relevant interface patches depending on the solvent-accessible surface area exposed by the ligand (Fig. 2B), and searched for complementary fingerprints in our seed database. Top-ranking seeds were selected, refined, and grafted onto recipient scaffolds, and approximately 2000 designs per target complex were selected with computational filters (Fig. 2C and table S1, see Methods). Our pipeline generated designs with diverse helical and beta sheet-based binding motifs, as well as various protein folds, thus sampling a wide space of sequences and topologies (Fig. 2C). All selected designs were predicted to favorably engage the neosurface by showing increased interface structural metrics in the presence of the ligand, such as the predicted binding energy, the buried surface area and the number of atomic contacts (fig. S3).

### Experimental validation of ligand-induced PPIs

The computational designs were screened by yeast display(*39*) and, after two rounds of fluorescent-activated cell sorting (FACS), enriched clones were deep sequenced (fig. S4 and table S2). We show one binder targeting each of the selected test cases (Fig. 3A). The best designs show no binding in the absence of the corresponding small molecules, whereas modest to high binding signals were observed with the ligands in yeast display experiments (Fig. 3B). These changes of binding signal upon small molecule addition are consistent with the expected behavior of a chemically induced PPI. Interestingly, small molecules contributed about 10-12% of the predicted target buried surface area, but they improved the predicted binding energy (ΔΔG) of the interface compared to the ligand unbound form by 17% to 27.7%. This result demonstrates a small, yet critical contribution that each ligand plays in the binding event, highlighting the difficulty of the design problem (table S3).

**Fig. 3:**
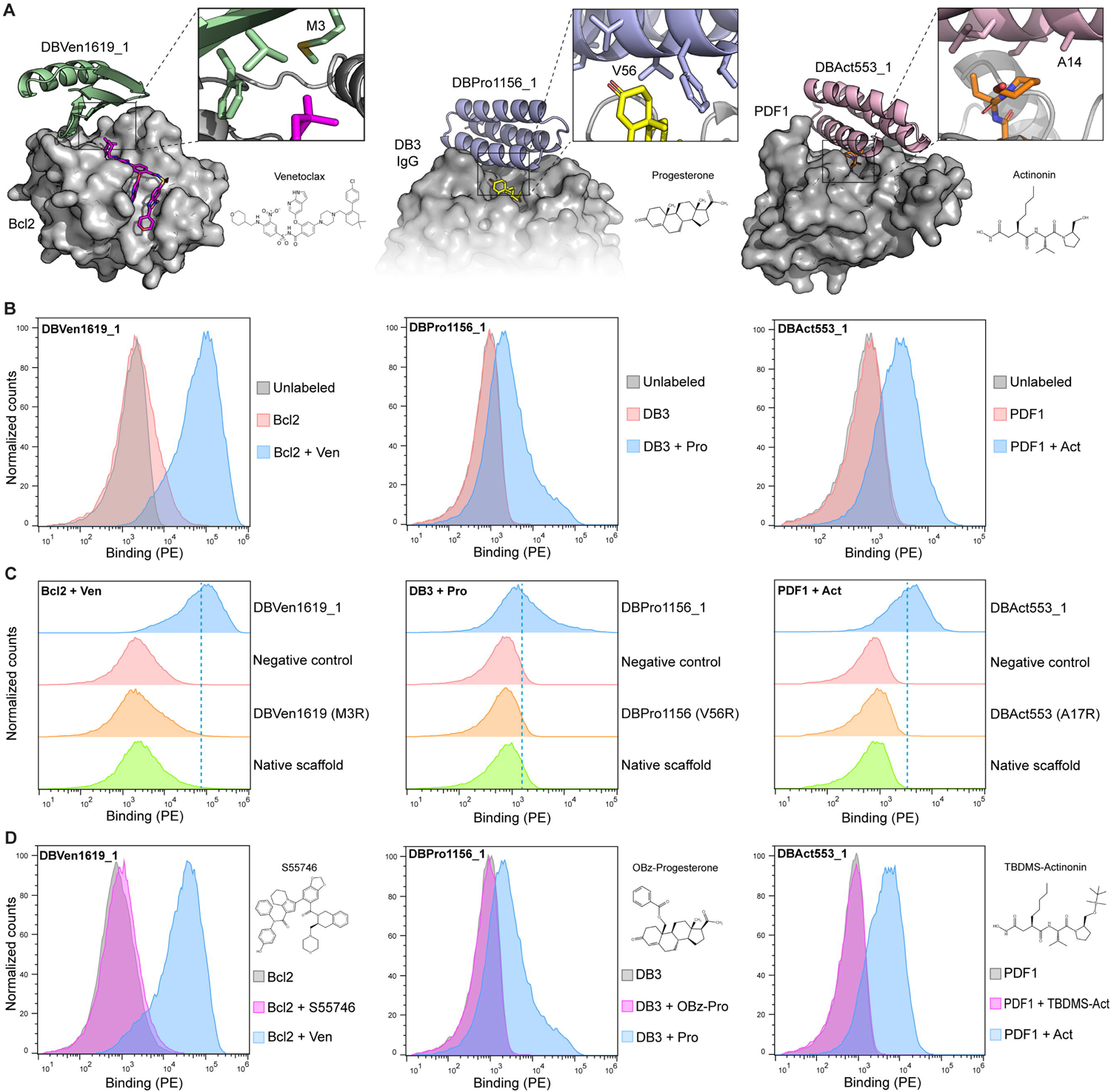
*De novo* design and screening of small molecule-dependent binders. **A.** Models of the designed binders in complex with their respective target complexes: Bcl2:Venetoclax, DB3:Progesterone and PDF1:Actinonin **B.** Histograms of the binding signal (PE, phycoerythrin) measured by flow cytometry on yeast displaying the designed binders. Yeast were either unlabeled or labeled with 500 nM of their respective target protein preincubated with the ligand, or with the target protein alone. **C.** Histograms of the binding signal (PE, phycoerythrin) measured by flow cytometry on yeast displaying designed binders, a mutated version with a single point mutant at the predicted interface and the starting scaffold used for the design process. Yeast cells were labeled with 500 nM of their respective drug:protein complex. Dashed lines represent the geometric mean of the designed binder signal. **D.** Binding measured on yeast displaying DBVen1619_1, DBPro1156_1 or DBAct553_1 labeled with the target protein alone (gray), the target protein in complex with the original small molecule (blue), or the target protein in complex with the small molecule analog (magenta). Control analogs tested were S55746, Progesterone-19-O-Benzoyl (OBz-Pro) and Tertbutyldimethylsilyl-Actinonin (TBDMS-Act).

Moreover, point mutants at the interface hotspot residues abrogated binding to the target complex, which further supports the designed binding mode (Fig. 3C). No binding was observed with the native scaffolds used for the seed grafting and interface design, underlying the critical role of the interface design pipeline (Fig. 3C). Finally, specificity towards the desired ligand was confirmed by using control compounds: S55746 for Bcl2, 19-O-Benzoyl-Progesterone (OBz-Pro) for DB3 IgG and Tertbutyldimethylsilyl-Actinonin (TBDMS-Act) for PDF1 (Fig. 3D and fig. S5). These analogs retained binding to the protein target (fig. S5). However, no binding to the designs was observed, confirming that the correct interface on the target complex is engaged with high ligand specificity (Fig. 3D).

### Biochemical characterization and structural validation

To map the binding site with high confidence and identify potential beneficial mutations (fig. S6), we performed a site-saturation mutagenesis (SSM) study (fig. S6). To assess the effect of the different mutations over the designed ligand-dependent interaction, we computed the average enrichment score of each mutation when comparing binding versus non-binding populations on yeast display experiments, similar to other deep saturation mutagenesis studies(*40*, *41*). Globally, we observed that such interactions have exquisite sensitivity to single-point mutants and that residues with high sensitivity mapped very closely to the designed interfaces, supporting the accuracy of our computational models (Fig. 4A).

**Fig. 4:**
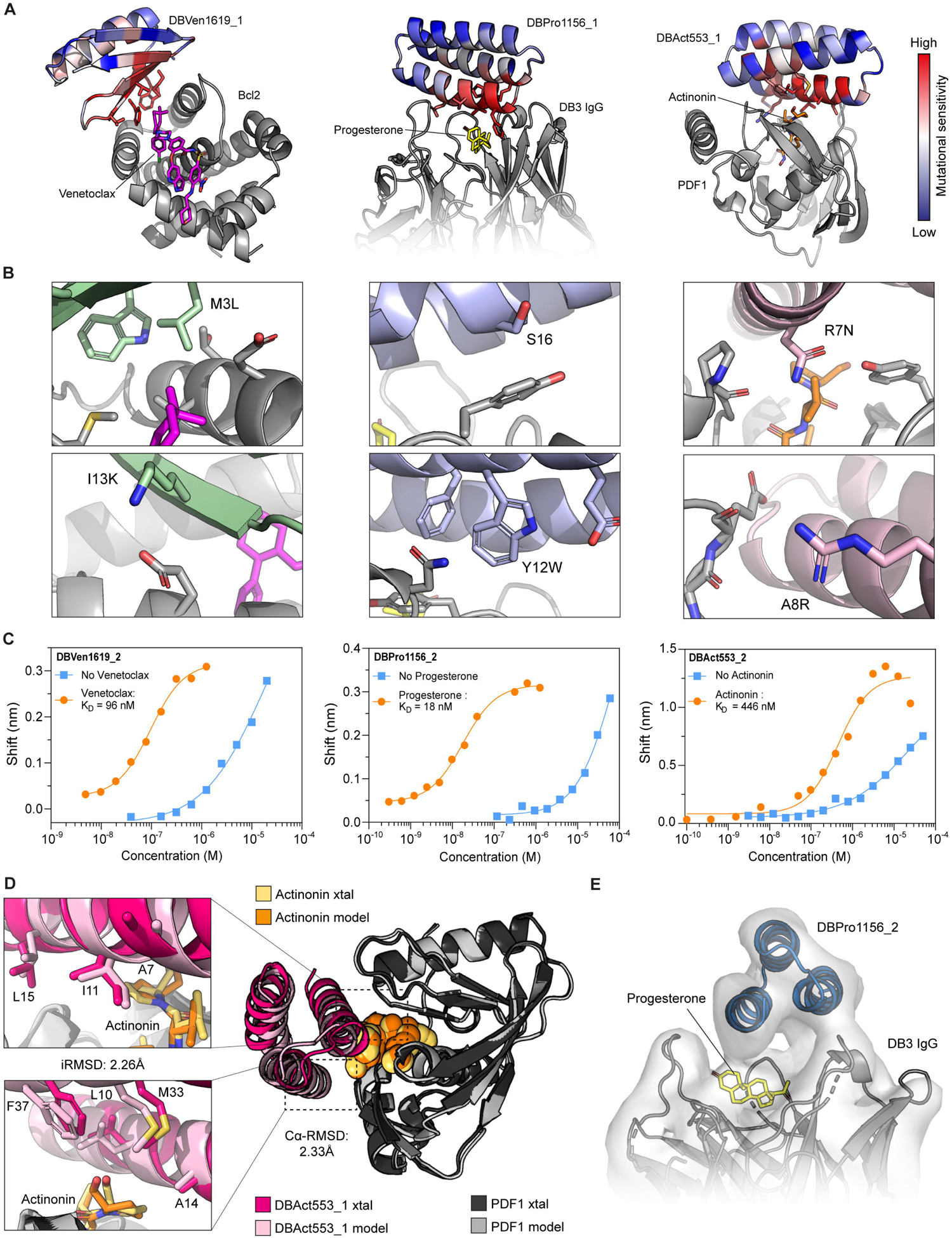
Optimization, characterization and functionalization of the designed binders. **A.** Computational model colored with the average enrichment score of the site-saturation mutagenesis of each amino acid position of the designed binder. Red color suggests that an amino acid position is sensitive to mutations, while blue color highlights a more tolerant amino acid position. Target proteins are shown in gray. **B.** Computational models incorporating the beneficial mutations that improved the affinity of the designed binders. Target proteins are shown in gray and designed binders in their respective color. **C.** Affinity measurement for DBVen1619_2, DBPro1156_2 and DBAct553_2 performed by biolayer interferometry. Each measurement was performed in presence (orange) or absence (blue) of the respective small molecule. The fits were calculated using a nonlinear four-parameter curve fitting analysis. **D.** Crystal (xtal) structure of DBAct553_1 in complex with Actinonin-bound PDF1 (PDB: 8S1X). The computational model (light pink) is aligned with the crystal structure (magenta). Inset: the alignment of the residues at the interface. E. Cryo-electron microscopy densities obtained for DBPro1156_2 in complex with progesterone-bound DB3.

The initial successful designs were expressed and purified for further biophysical characterization. All designs were monomeric, folded and highly stable in solution (fig. S7). All three designs showed binding affinities in the range of native transient PPIs(*42*), from mid-nanomolar to low-micromolar, after pure *in silico* generation (fig. S8). Specifically, DBAct553_1 showed a binding affinity (K_D_) of 542 nM, DBVen1619_1 and DBPro1156_1 showed affinities of 4 μM and >10 μM, respectively.

In the SSM scan, some mutations suggested potential improvements in affinity (fig. S6). Due to the large number of beneficial mutation candidates for DBVen1619_1, we created a combinatorial library covering 6 residues, sampling a set of favorable amino acids identified by SSM (fig. S9). Three of the six positions converged into single mutations (K1Q, M3L, I13K) while the remaining three residues did not converge. We engineered a variant, DBVen1619_2, with the three beneficial mutations and confirmed the binding improvement on yeast display (fig. S9). Among the favorable mutations, M3L in the core of the interface between Bcl2:Venetoclax and DBVen1619_2 plays a crucial role (Fig. 4B). The conformational rigidity of a leucine is likely to be preferred to the rotameric flexibility of a methionine(*43*), reducing the entropic cost of the binding interaction(*44*). On the other hand, the second beneficial mutation (I13K) is likely to provide a favorable electrostatic interaction with a glutamate nearby. Overall, the incorporation of the three mutations resulted in a 42-fold improvement of the affinity (K_D_ = 96 nM, Fig. 4C)

For the progesterone-dependent binder, DBPro1156_1, four favorable mutations were identified by SSM and showed an increased binding on yeast display (fig. S10). Two mutations (Y12W and S16G) significantly improved the binding signal and showed an additive effect in the resulting design, DBPro1156_2. Modeling of the two mutations suggested increased interface packing (Y12W) and the removal of a steric clash (S16G) (Fig. 4B, middle panel). DBPro1156_2 showed a binding affinity of 18 nM, which represents an improvement of three orders of magnitude, relative to the parent design, solely with two mutations (Fig. 4C).

Several mutations were found to slightly improve binding of DBAct553_1 to Actinonin-bound PDF1 (fig. S11). Most of these mutations were hypothesized to result in a more elaborate hydrogen bond network across the interface (e.g. R7N or A8R) (Fig. 4B). Of note, the combination of I3E with R7N was found to be deleterious for binding (fig. S11), most probably because of their spatial proximity that might trigger unwanted side chain rearrangement. A combination of the beneficial mutations (R7N and A8R) gave rise to DBAct553_2, which bound with an affinity of 446 nM for the Actinonin-bound PDF1 (Fig. 4C).

To evaluate the structural accuracy of our computational design approach, we co-crystalized the ternary complex of Actinonin-bound PDF1 with DBAct553_1 (PDB: 8S1X, Fig. 4D). The crystal structure closely resembled the computational model with a C_α_ RMSD (Root Mean Square Deviation) of 2.33 Å and a full-atom interface RMSD (iRMSD) of 2.26 Å, which demonstrates the accuracy of our design pipeline. The deviation from our initial model can to a large extent be attributed to a misplaced residue (Y2) in the model of the design scaffold which induced a slight shift of the N-terminal helix (fig. S12). Consequently, the C_α_ RMSD of our model deviates 0.93 Å from that of the experimental structure (fig. S12). Of note, the AlphaFold2(*45*) prediction of the monomeric designed binder aligned perfectly with our structure with a C_α_ RMSD of 0.49 Å by placing residue Y2 with the correct orientation. Overall, this observation together with previous findings suggests that an increased use of deep learning tools like AlphaFold should significantly increase the model accuracy and therefore success rate(*46*). Finally, we obtained low-resolution cryo-electron microscopy densities of the DBPro1156_2 in complex with DB3 Fab and progesterone that confirmed the designed binding mode and interface engagement with the small molecule (Fig. 4E). Despite the absence of structural data for the remainder of the designs, the mutational sensitivity assessed by the SSM (Fig. 4A) and the lack of binding with the small molecule analogs (Fig. 3D) suggests that the binders engage the target interface with a binding mode in agreement with our computational models.

### Designed CIDs are functional in cell-based systems

Chemically controllable components have important applications in synthetic biology and have been shown to be useful in modulating the activity of emerging cell-based therapies (*10*, *11*, *47*). To test whether our computationally designed CIDs would assemble in a more complex cellular context, we engineered reporter proximity-based systems that were expressed in cell-free system or mammalian cells and that in the presence of the small molecule could activate a signaling pathway or lead to the reconstitution of a reporter protein. The most natural functional logic for chemically-induced protein interactions is to function as ON-switch systems.

We first repurposed a previously described heterodimerization-based reporter system(*48*) to test the DB3 antibody as a single-chain variable fragment (scFv) binding to DBPro1156_2. Here, DB3 was fused to a zinc finger 438 transcription factor and DBAct553_2 to a T7 RNA polymerase (Fig. 5A), and tested in a cell-free reporter system. The heterodimerization in presence of the drug induces proximity between the T7 RNA polymerase and the transcription factor, thus leading to the transcription of a reporter linear DNA template and its translation into a red fluorescent protein (mCherry). While only baseline fluorescence was observed in absence of progesterone, a 15.8-fold increase was observed after addition of progesterone (Fig. 5B). Similarly, a titration of progesterone demonstrated a dose-response curve, suggesting possible utilization as a novel cell-free biosensor (Fig. 5C).

**Fig. 5:**
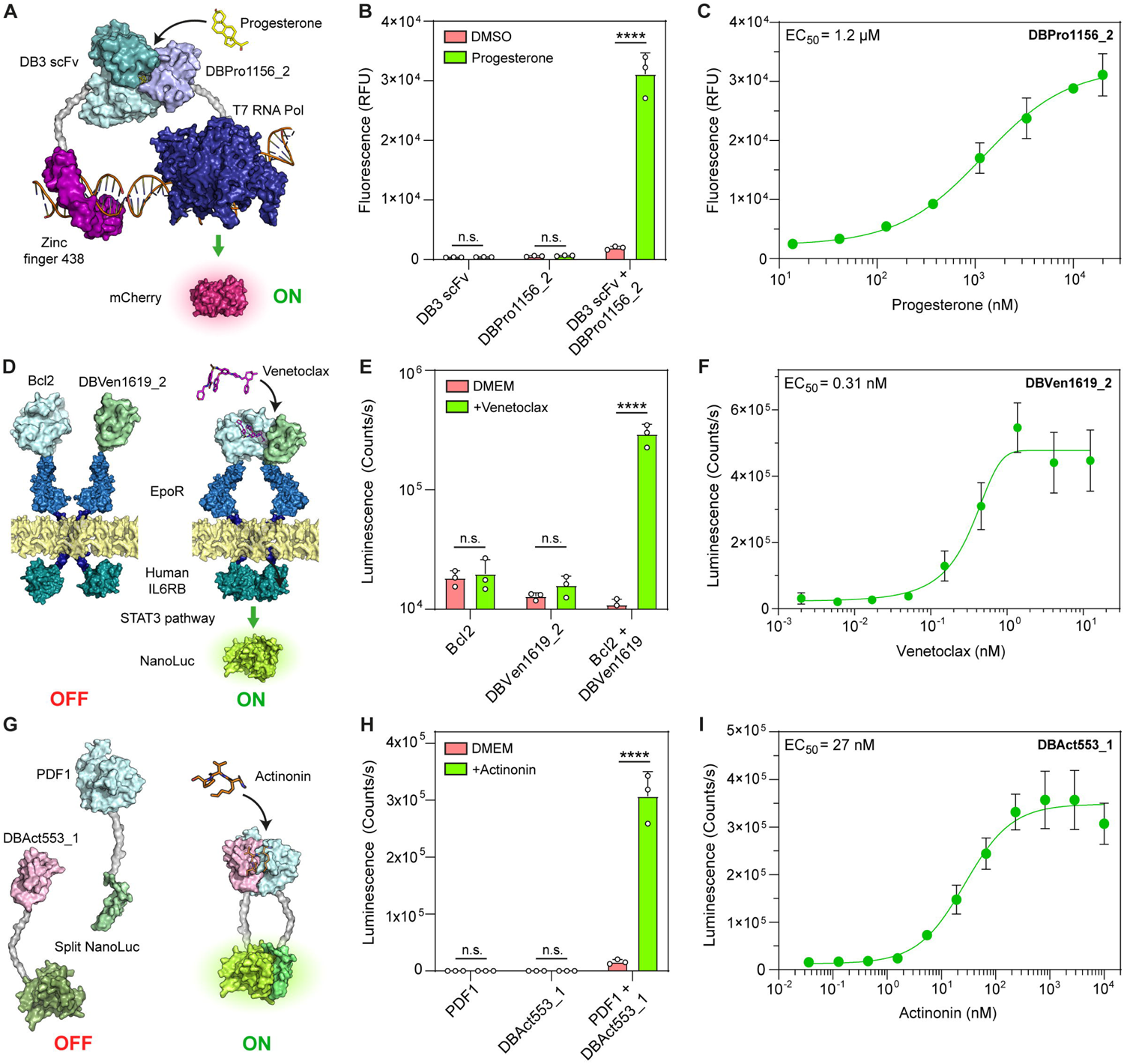
Computationally designed CIDs are functional in cell-based systems. **A.** Schematic of the cell free-expression system with single chain variable fragment (scFv) DB3-fused to a zinc finger transcription factor and DBPro1156_2 fused to T7 RNA polymerase. **B.** Fluorescence (Relative fluorescence unit; RFU) measured in wells containing each monomeric component or mixed, without or with 20 μM progesterone. **C.** Progesterone dose-dependent responses performed in a cell free system containing both components. **D.** Schematic of the GEMS reporter system functionalizing Bcl2-based CID. Both protein components of the CID are individually fused to erythropoietin receptor (EpoR) chains linked to an intracellular human IL6RB domain, which induces the expression of a reporter gene (secreted NanoLuc luciferase) when activated. **E.** NanoLuc luminescence of HEK293 cells transfected with Bcl2-GEMS only, DBVen1619_2 only or both together without or with 1 μM Venetoclax. **F.** Venetoclax dose-dependent responses performed on HEK293 transfected with Bcl2 and DBV1619 GEMS receptors. **G.** Schematic of the split NanoLuc system functionalizing DBAct553_1 and PDF1. **H.** Intracellular NanoLuc luminescence of HEK293 transfected with C-term split NanoLuc-fused PDF1 only, N-term split NanoLuc-fused DBAct553_1 only or both together without or with 25μM Actinonin. **I.** Actinonin dose-dependent responses performed on HEK293 transfected with split-NanoLuc PDF1 and DBAct553_1. p < 0.0001 (****), non-significant (ns).

To test the chemically-induced activity of the designed modules in mammalian cells, we used a previously described system called generalized extracellular molecule sensor (GEMS)(*13*). Briefly, the target protein and the designed binder are both fused to an erythropoietin receptor (EpoR) linked to an intracellular domain of a human interleukin 6 receptor subunit B (IL6RB) (Fig. 5D). Transcription of a reporter gene (NanoLuc luciferase)(*49*) will be triggered upon a conformational change induced by the heterodimerization in presence of the drug. By incorporating Bcl2 and DBVen1619_2 in the GEMS system, we observed a 26.8-fold change in luminescence in the presence of Venetoclax, while minimal background was observed in the absence of the drug (Fig. 5E). These results show the desired behavior of an ON-switch system. Additionally, our modified GEMS system displayed a heightened sensitivity to the drug, with a half maximal effective concentration (EC_50_) of 0.31 nM, which is likely due to the co-localization of the sensing modules in the cell membrane (Fig. 5F).

Next, we designed a cytoplasmic system to respond to Actinonin and fused PDF1 and DBAct553_1 to two moieties of a split NanoLuc (Fig. 5G). In this system we also observed a significant increase in signal (19.1-fold) upon dosing of the cells with Actinonin (Fig. 5H). This novel ON-switch system was also highly sensitive to the presence of the drug, as shown by the titration reporting an EC_50_ of 27 nM (Fig. 5I). Overall, we showed that our computationally designed CIDs can be used to functionalize molecular components in cellular systems, suggesting a promising route for the development of new modules for synthetic biology including a wide range of biosensors and cell-based applications.

## Discussion

Current deep learning-based protein design pipelines are primarily conditioned on the natural amino acid repertoire(*50–52*) and therefore lack generalization to the design of interactions involving small molecules. This gap is mainly due to the scarcity of protein-ligand structural data, and especially ternary complexes, within the training sets based on the PDB, where such complexes are limited(*53–55*). Geometric deep learning approaches principled in the physical and chemical features of the molecular surface can overcome these limitations, and provide joint representations for protein and small molecule complexes. The resulting neosurfaces capture and present generalizable molecular features that enable the challenging task of designing protein binders targeting these hybrid interfaces. Utilizing the *MaSIF-neosurf* framework, we successfully designed three specific binders against Bcl2:Venetoclax, DB3:Progesterone and PDF1:Actinonin complexes. All designed binders showed high stability, specificity and native-like affinity for their target complexes by pure *in silico* generation. The affinities were experimentally optimized to nanomolar range and their binding mode was confirmed through mutational and structural characterization, showcasing the accuracy of our design pipeline. Notably, our pipeline managed to capture the subtle, yet crucial contributions of each ligand (10-12% of the buried SASA only; table S3) to induce protein interactions. This sensitivity represents an additional layer of complexity to the task of designing highly sensitive CIDs, compared to previous attempts targeting large ligand interfaces(*26*).

To demonstrate the functionality of our designed CID systems, we probed their efficiency and specificity in the context of a complex cellular environment. They exhibited robust ON-switch behavior in both cytoplasmic and membrane-bound circuits, showcasing their potentially wide applicability in mammalian systems as logic gates, synthetic circuits, or new biosensors for detecting specific metabolites(*10*, *13*). This relevance is further underscored by our use of the FDA-approved drug Venetoclax for treating leukemia(*29*) the natural product Actinonin with potentially chemotherapeutic effects (*31*) or the endogenous hormone progesterone (*56*). These can be utilized for combined anti-cancer therapies with chimeric antigen receptor (CAR) T cells, which are often hindered by off-target toxicities (*11*, *57*). The addition of synthetic small molecule activators could allow finer control of their activity and elevate their safety profile.

While the design of specific protein-ligand interactions remains challenging, the presented results lay a strong foundation for further innovations. Incorporation of deep learning-based structure validation methods, such as AlphaFold2(*45*) (fig. S13) or RoseTTAFold(*58*), or generative models complemented with surface fingerprints(*51*) could improve design success rates. Additionally, accounting for conformational flexibility and dynamics at the surface could pave the way for more complex interaction types, such as intrinsically disordered proteins. Overall, we envision that surface-based representation can contribute to solving molecular design problems in low-data regimens, such as the design of protein-based molecules with non-natural amino acids. The capability of extracting expressive fingerprints from protein:ligand complexes opens up the tantalizing possibility of rationally designing innovative drug modalities, such as on-command cell-based therapies(*11*, *17*), controllable biologics(*21*, *23*), or molecular glues, which thus far remains an outstanding challenge in drug development(*8*, *9*)

## Acknowledgments

We would like to thank the staff at PTPSP at EPFL, Florence Pojer, Kelvin Lau, Amédé Larabi, Laurence Durrer and Soraya Quinche for their advice on the biophysical characterization of proteins and their work for the structural validations; the staff at the Dubochet Center for Imaging (DCI) in Lausanne for cryo-EM data collection and processing; SCITAS at EPFL for support in the computational simulations; the staff at GECF for assistance with deep sequencing and members of FCCS for assistance in FACS. We also thank Dr. Andrea Moretti from Malvern Panalytical for his support to run the Grating-Coupled Interferometry on Creoptix Wave and giving access to the instrument. We express our gratitude to Prof. Eva-Maria Strauss for providing *de novo* hyperstable proteins for our scaffold database. Finally, we thank Prof. Nicolas Thomä and Dr. Kelvin Lau for their feedback on the manuscript.

## Author contributions

A.M., S.B. and A.S. contributed equally to this work. A.M. and B.E.C led the project. A.M., S.B., P.W.L. and M.E. performed the experimental work. A.M., S.B., M.P. and L.S. designed the experimental methodology. A.M. and A.S. performed the computational work and protein design. A.M., A.S., P.G., E.E. and R.M.N. contributed to the design of the computational pipeline. M.P. solved the crystal structure. L.R. synthesized the small molecule analogs. S.G. and J.S. participated in the expression and purification of proteins. P.S., S.J.M., M.B. and B.E.C provided supervision and acquired the necessary funding. A.M., S.B., A.S., M.P. and B.E.C wrote the manuscript with inputs from all authors.

## Data and material availability

Crystal structure of DBAct553_2 in complex with Actinonin-bound PDF1 has been deposited at the PDB under the accession code 8S1X (DOI: https://doi.org/10.2210/pdb8S1X/pdb). *MaSIF-neosurf* and the Rosetta design scripts are available on GitHub (https://github.com/LPDI-EPFL/masif-neosurf). The scaffold database generated for grafting the seed provided by *MaSIF-neosurf* is partly available at Zenodo (https://zenodo.org/records/7643697#.Y-z533ZKhaQ) and partly on Github (https://github.com/strauchlab/DBP and https://github.com/strauchlab/scaffold_design/). All other data needed to evaluate the conclusions in this paper are present either in the main text or the supplementary materials.

