## Supplementary materials for "Targeting protein-ligand neosurfaces using a generalizable deep learning approach"

### Materials and methods

Incorporation of small molecules in *MaSIF-seed*

Molecular surface meshes were triangulated using the MSMS program(*59*) and radial patches (geodesic radius 12Å) computed following the original MaSIF preprocessing scripts(*27*). Before applying MaSIF's geodesic convolutional layers, five input features are computed for each patch: shape index(*32*), distance-dependent curvature(*33*), Poisson-Boltzmann continuum electrostatics, hydrogen bond donor and acceptor potential(*34*), and hydropathy(*35–37*). The first two features are purely geometric and are calculated analogously to protein surfaces alone. Moreover, the APBS program(*60*) used for computing the Poisson-Boltzmann electrostatics on the surface supports small molecules in the MOL2 file format and hence does not require us to treat them in a conceptually different way. The remaining two chemical input features are computed as described below.

*Hydrogen bond donors and acceptors*

The hydrogen bond propensity feature assigns a positive value to points on the molecular surface near the optimal direction in which a hydrogen could be formed with an acceptor atom. It is determined by the direction of the covalent bond between a donor atom and its hydrogen (fig. S1B-C). Likewise, a negative value is assigned to points corresponding to hydrogen bond acceptors. For different acceptor types, the theoretically optimal position for forming a hydrogen bond can either lie on a cone (fig. S1D-F) or in a small number of specific directions that can be derived from the molecular geometry. We assign different magnitudes of the donor/acceptor feature based on the angular deviation from the ideal hydrogen bond geometry according to a quadratic function.

The optimal direction of the hydrogen bond is determined using the RDKit software package(*61*) and surface points are assigned positive (donor) or negative (acceptor) values between -1 and +1 based on their angular deviation from the ideal direction. For potential acceptors, RDKit was also used to determine whether the idealized location of the hydrogen bond lies on a cone or in one or more discrete directions.

*Hydropathy*

MaSIF's hydrophobicity feature makes use of the Kyte-Doolittle scale(*35*) which is exclusively defined for amino acids. Equivalent values for small molecules thus need to be approximated based on a more general hydrophobicity measure that can be estimated computationally, such as the logarithm of the octanol−water partition coefficient (LogP)(*36*). To this end, we develop a nonlinear function that maps LogP values to the KD scale. We fit the parameters of this function to find an optimal match for the KD and LogP values of all twenty amino acids. Since the best functional form of this mapping is not immediately obvious from the raw values (fig. S1L), we experimented with different hydrophobicity scales as intermediates and found that the Eisenberg scale(*37*) has approximately linear and exponential relationships with LogP and KD-values of amino acids, respectively. We first compute the optimal parameters of the mappings from LogP to Eisenberg scale (fig. S1G) and Eisenberg scale to Kyte-Doolittle scale (fig. S1H) and then compose these two functions to establish the desired relationship between LogP and KD values (fig. S1H). Finally, we also restrict the outputs to the valid interval of KD values [-4.5, 4.5] to ensure that the feature does not leave the domain MaSIF was trained on.

Furthermore, since some ligands can cover large surface patches, we aim to respect local variations of the hydrophobicity by fragmenting the molecules prior to calculating their hydrophobicity score. We employ the BRICS algorithm(*62*) to decompose molecules, and compute estimates of each fragment's logP value with RDKit. The resulting fragments are more similar in size to amino acids and tend to have less extreme hydrophobicity scores than whole ligands, moving the distribution of this feature closer to that expected on protein surfaces (fig. S1K-L). To translate from logP to the Kyte-Doolittle scale, we parameterize a function so that it approximates the relationship between these hydrophobicity values for the 20 amino acids. KD and Eisenberg values of all amino acids are available in tabular form, whereas we compute their LogP with RDKit to fit the curves. The final function is

$$KD = clip\,(-6.2786 + exp\,(0.4772 * logP + 1.8491), min = -4.5, max = 4.5).$$

After computing equivalent KD values for all small molecule fragments, we assign the resulting hydrophobicity score of the closest fragment to each surface vertex.

To create the histograms in fig. S1G, we extracted 20,363 unique small molecule ligands from the Binding MOAD(*63*) dataset, fragmented each, and removed duplicates. This resulted in 9,362 unique fragments that are compared to the set of ligands and the twenty standard amino acids.

<u>Binding site identification</u>

MaSIF-site(*27*) was trained on a dataset of known PPIs to predict regions on protein surfaces with high propensity for forming a buried interface. The neural network takes a protein-ligand complex decomposed into 12 Å (geodesic radius) overlapping patches as input and generates a per-vertex regression score, indicating the propensity of each point to become a buried surface area within a protein interaction. In this study, we employed MaSIF-site to predict interfaces and guide the selection of target patches both in our computational benchmark and for all three target complexes for design (Bcl2:Venetoclax, DB3:Progesterone and PDF1:Actinonin). In the computational benchmark, we conducted the search only for the three patches with the highest interface propensity near the center of the binding site. For design, the number of targeted sites overlapping with the protein-ligand neosurface depended on the solvent-accessible surface area of each ligand to ensure that all the ligand-exposed surface was covered during the complementary motif search: 1 for PDF1:Actinonin, 2 for DB3:Progesterone and 3 for Bcl2:Venetoclax.

Binding seed identification

The fingerprints of the predicted 12 Å (geodesic radius) patches comprising both protein target and bound drug were used to find a complementary fingerprint in the MaSIF-seed database(*28*) which consists of ~640’000 continuous structural fragments (seeds) amounting to 402 million surface patches/fingerprints. The seed database covers distinct secondary structures with approximately 390’000 sheet-based and 250’000 helical motifs respectively. The MaSIF-search algorithm was trained to make patch fingerprints similar for interacting patches and dissimilar for non-interacting patches. Seeds with interface propensity scores above the defined threshold and with fingerprint distances (Euclidean distance between target and seed fingerprint) below the defined thresholds were selected. In a second-stage alignment and scoring using the RANSAC algorithm, seeds were selected based on interface post-alignment (IPA) score. Cutoffs used for the seed selection are summarized in table S1.

Scoring aligned structures

We consider two descriptor-based post-alignment scores. The descriptor distance score (DDS) is a simple heuristic that aggregates descriptor distances across the predicted binding interface. DDS is based on the squared Euclidean distances between interacting patches on both sides of the interface. Two patches are considered interacting with each other if their center points are less than 1.5Å apart. The descriptor distance score is computed according to the following formula

$$DDS = \sum_{i} \frac{1}{||\, binder\_desc(i) - target\_desc(NN(i))\, ||^2}$$

where *i* indexes interacting patches of the first protein and *NN(i)* returns the index of the spatially nearest neighbor on the other protein. Higher scores mean higher complementarity.

The interface post-alignment score (IPA) is computed by a neural network that was trained to discriminate between near-native and high-RMSD poses of docked proteins(*27*). The inputs of this predictor are 3D Euclidean distances, descriptor distances and dot products between surface normals of up to 200 pairs of corresponding patches at the predicted interface. It outputs values between zero and one where larger values indicate higher confidence in the presented interface.

Computational binder recovery benchmark

The binder recovery experiment was performed for 14 known ligand-induced protein complexes, where both proteins involved in the interaction are considered as separate items, resulting in 28 search queries. Additionally, we included 200 decoys (based on 100 PPIs) in the database. All benchmark complexes and decoys are listed in table S4. After triangulating and featurizing all protein surfaces with and without ligands, we screen the database and dock candidates analogously to the binding seed search. Here, we assume the location of the binding site on the target protein is known and select the three surface vertices with the largest predicted surface propensity within 10Å of the center of this site as input patches. The center of the binding site was approximated with a simple heuristic. We first identify interface atoms as those within 4Å of any atom from the binding protein in the original complex structure. This can and typically will include atoms belonging to the small molecule. Then we define the average of the coordinates of all interface atoms of the target protein as the center of the binding site. Furthermore, we declare a binder correctly recovered if its interface RMSD (iRMSD) compared to the ground truth structure of the same protein is less than 5Å, where iRMSD considers only heavy atoms in the immediate vicinity of the target protein (<5Å).

Seed and interface refinement

To optimize binding energy of the seed for the target complex, seeds were refined using a FastDesign protocol on Rosetta(*38*) with a penalty for buried unsatisfied polar atoms in the scoring function(*64*). Refined seeds were then selected based on the computed binding energy (ddG), shape complementarity, number of interface hydrogen bonds, number of buried unsatisfied polar atoms and number of atoms in contact with the small molecule. Beta sheet-based motifs making >33% contact with the target complex using loop regions were discarded. Moreover, uniqueness of each seed was assessed by doing a pairwise alignment of the hotspot residues. For seeds showing >70% hotspot identity with another seed, only the one with the best surface-normalized ddG was kept.

Seed grafting and computational design

Selected seeds were subsequently grafted with a Rosetta MotifGraft(*65*) protocol for stabilizing the binding motif and bringing additional contacts with the target complex. Each seed was match with a database of ~6500 small protein scaffolds (<90 amino acids) originating from small globular monomeric protein from the protein data bank (PDB)(*66*) and four computationally designed miniprotein databases that were experimentally validated(*67–70*). Prior grafting, seeds were cropped to the minimum number of residues making contact with the target, and loop motifs were removed from beta sheet-based seeds for optimizing the grafting success rate. Once grafting was performed, scaffolds underwent sequence optimization using a FastDesign protocol on Rosetta with a penalty for buried unsatisfied polar atoms in the scoring function. Final designs were selected based on the computed binding energy (ddG), shape complementarity, number of interface hydrogen bonds and count of buried unsatisfied polar atoms. A similar number of designs per seed was ensured by setting dynamic cutoffs of these metrics adjusted for each seed.

Library screening

For each target complex, around ~2000 designs were reverse-translated into DNA and purchased from Twist Bioscience as oligo pools with 18bp homology overhangs. Oligo pools underwent two rounds of PCR : i) for the amplification of the library using the 18bp overhangs and ii) for adding 45bp homology with the yeast display vector (57.5 °C annealing for 30 s, 72 °C extension time for 1 min, 15 cycles). EBY-100 yeast were transformed by electroporation using the amplified inserts and linearized HA-tagged pCTcon2 vector as described previously(*39*). A similar approach was done for site-saturation mutagenesis (SSM) library of single designs. Transformed yeast cells were grown in minimal glucose medium (SDCAA) medium at 30°C and induced with minimal galactose medium (SGCAA) medium overnight prior sorting.

Yeast surface display of single designs

Genes encoding for single designs were purchased from Twist Bioscience with a ~25bp homology overhang for cloning. Each design was cloned into HA-tagged pCTcon2 plasmid using Gibson assembly and transformed into XL10-Gold or HB101 bacteria for DNA production. The purified and sequence-approved DNA was then used to transform competent EBY-100 yeast using the Frozen-EZ Yeast Transformation II Kit (Zymo Research). As for libraries, transformed yeast cells were grown in minimal glucose medium (SDCAA) medium at 30°C and induced with minimal galactose medium (SGCAA) medium overnight prior flow cytometry analysis.

Flow cytometry analysis and sorting

Induced yeast cells were washed with PBS supplemented with 0.1% BSA and then labeled with the respective binding target for 2 hours at 4°C. Prior to labeling, protein-drug complexes were pre-incubated at room temperature for 5 min with a 1:5-10 ratio. Cells were then washed and labeled with a FITC-conjugated goat anti-HA tag antibody (Bethyl; ref: A190-138F; display tag; 1:100 dilution) and a PE-conjugated goat anti-human Fc antibody (Invitrogen; ref: 12-4317-87; binding tag; 1:100 dilution) for 30 min at 4°C. Cells were washed, resuspended in an appropriate volume of buffer and analyzed on a Gallios flow cytometer (Beckman Coulter), or sorted with a Sony SH800 cell sorter. Kaluza software (Beckman Coulter, v.1.1.20388.18228) and LE-SH800SZFCPL Cell Sorter (Sony, v.2.1.5) were respectively used for the data acquisition. In the case of cell sorting, each designed library was sorted for binding and non-binding populations separately. Flow cytometry data were then analyzed using FlowJo (BD Biosciences, v.10.8.1).

Library sequencing

Sorted yeast were cultured and plasmids encoding protein designs were extracted using the Zymoprep Yeast Plasmid Miniprep II (Zymo Research) following the manufacturer's protocol.The sequence of interest was then amplified by PCR with vector-specific primers flanking the protein design gene. A second PCR was performed to add Illumina adapters and Nextera barcodes, and the PCR product was desalted and purified using the Qiaquick PCR purification kit (Qiagen). Illumina MiSeq system with 500 cycles was used for the next generation sequencing. Around 0.8-1.2 millions reads per sample were obtained, translated into the appropriate reading frame and matched with expected input sequences from the libraries. The enrichment of each design was calculated by normalizing the counts in the binding population with the counts in the non-binding populations. Hits were identified if the enrichment was >10-fold and the number of counts in the binding population was >10'000.

Protein expression and purification

A list of protein sequences can be found in table S5. Genes encoding the 6xHis-tagged and/or human Fc-tagged protein of interest were purchased to Twist Bioscience cloned into pET11 (bacteria vector) or pHLSec (mammalian vector) by Gibson assembly and transformed into XL10-Gold or HB101 bacteria. Plasmids were extracted using a GeneJET plasmid Miniprep kit (Thermofisher, for bacteria vector) or a PureLink Fast Low-Endotoxin Midi plasmid purification kit (Invitrogen, for mammalian vector) and checked by Sanger sequencing. Proteins were purified by bacteria or mammalian expression systems. Mammalian expressions were performed using the Expi293 expression system (ThermoFisher; ref: A14635). Supernatants were collected after 6 days, filtered and purified as explained below. For bacteria expression, BL21(DE3) or T7 Express Competent E. coli were transformed with the plasmid of interest and grown as a pre-culture overnight. Pre-cultures were inoculated 1:50 in Terrific Broth medium and incubated at 37°C until they reached a density ~0.7 at $OD_{600}$. Then, bacteria were induced with 1mM IPTG and incubated overnight at 18-20°C. Cells were collected by centrifugation at 4000g for 10 min, resuspended in lysis buffer (50 mM Tris, pH 7.5, 500 mM NaCl, 5% glycerol, 1 mg $ml^{-1}$ lysozyme, 1 mM PMSF and 1 µg $ml^{-1}$ DNase) and lysed by sonication. Lysates were then clarified by centrifugation at 30'000g for 30 min and filtered.

All 6xHis-tagged protein are purified using the ÄKTA pure system (GE healthcare) Ni-NTA HisTrap affinity column followed by a size exclusion chromatography on a Superdex HiLoad 16/600 75pg or 200pg depending on the size of the protein. All proteins were concentrated in PBS as a final buffer.

Surface plasmon resonance

Affinity measurements were done on a Biacore 8K (GE Healthcare) using HBS-EP+ as a running buffer (10 mM HEPES at pH 7.4, 150 mM NaCl, 3 mM EDTA, 0.005% v/v Surfactant P20, GE Healthcare). All proteins were immobilized on a CM5 chip (GE Healthcare #29104988) via amine coupling to reach 500–1000 response units (RU). Analytes were then injected in serial dilutions using the running buffer. The flow rate was 30 µL/min for a contact time of 120 s followed by 400 s of dissociation time. SPR data were either fit with a 1:1 Langmuir binding model within the Biacore 8K analysis software (GE Healthcare #29310604) or done in steady-state affinity mode by reporting the relative RU for each concentration.

Biolayer interferometry

Biolayer Interferometry (BLI) measurements were performed on the Gator BLI system using the GatorOne software (Gator Bio, v.2.7.3.0728). Running buffer consisted of 500mM NaCl and 50mM Tris pH 7.5 or HPS-P+ buffer (10 mM HEPES pH 7.4, 150 mM NaCl, 1µM $NiSO_4$, 0.005% v/v Surfactant P20, GE Healthcare), supplemented by 100nM Venetoclax or 5µM Actinonin if needed. Fc-tagged proteins were immobilized at a concentration of 7µg/ml on protein A probes (1.5 to 2.5nm immobilized) and dipped into serial dilutions of the ligand. Steady state responses were normalized with the maximum value and plotted using a nonlinear four-parameter curve fitting analysis.

Grating-Coupled Interferometry

Grating-Coupled Interferometry (GCI) measurements were performed on a Creoptix WAVE system (Malvern Panalytical) using the Creoptix WAVE control software (Malvern Panalytical, v. 4.5.18). Running buffer consisted of HPS-P+ buffer (10 mM HEPES pH 7.4, 150 mM NaCl, 0.005% v/v Surfactant P20, GE Healthcare). All protein targets were immobilized on a 4PCH chip (Malvern Panalytical) via amine coupling to reach 7'000-10'000 pg/$mm^2$ . An intermediate injection with 1µM $NiSO_4$ was used for PDF1 protein. S55746, 19-O-Benzoyl-Progesterone (OBz-Progesterone) and Tertbutyldimethylsilyl-Actinonin (TBDMS-Actinonin) were then injected sequentially as analytes at a concentration of 2, 2.5 and 5 µM respectively using the waveRAPID (Repeated Analyte Pulses of Increasing Duration) kinetic assay(*71*). The flow rate was 100 µL/min for an injection duration of 25s followed by 300s of dissociation time for TBDMS-Actinonin, while an injection duration of 50s followed by 600s of dissociation time was used for S55746 and OBz-Progesterone. Measurements were either fitted with a 1:1 model for Bcl2:S55746 and PDF1:TBDMS-Actinonin, or with a mass transport model for BD3:OBz-Progesterone.

Size-exclusion chromatography–multi-angle light scattering

Size exclusion chromatography combined to multiangle light scattering device (miniDAWN TREOS, Wyatt) was performed to determine the molecular weight of the purified designs. The final concentration was approximately 1 mg/ml in PBS (pH 7.4), and 100 µl of the sample was injected into a Superdex 75 10/300 GL column (GE Healthcare) with a flow rate of 0.5 ml/min. $UV_{280}$, refractive index (dRI) and light scattering signals were recorded. Molecular weight was determined using the ASTRA software (version 6.1, Wyatt).

Circular dichroism

Far-ultraviolet circular dichroism spectra were carried with a Chirascan spectrometer (AppliedPhotophysics). Protein samples were prepared diluted in PBS at a protein concentration 300 µg/ml and placed in 1 mm path-length cuvette. Wavelengths between 200 nm and 250 nm were recorded with a scanning speed of 20 nm min−1 and a response time of 0.125 s. All spectra were corrected for buffer absorption. Temperature ramping melts were performed from 20 to 90 °C with an increment of 2 °C $min^{-1}$. Thermal denaturation curves were plotted by the change of ellipticity at the global curve minimum. If possible, melting temperature ($T_m$) were determined after fitting the data with a sigmoid curve equation on GraphPad Prism.

Cell transfection and induction

Human Embryonic Kidney (HEK293T; RRID: CVCL_0063) cells were cultured in Dulbecco's Modified Eagle Medium (41966-029, Gibco) supplemented with 10% (v/v) FBS (A5256701, Gibco) and 1%(v/v) antibiotic penicillin/streptomycin (15140-122, Gibco). Cells were maintained at 37°C with 5% CO2 and passaged every two to three days at around 80% confluency. Cells were seeded into the inner 60 wells of a 96 well plate, at 10'000 cells per well, 24 hours prior to transfection. Cells were transfected by layering 50uL from a mixture of 330uL DMEM, 825ng-850ng total DNA, and 4.125ug PEI (24765-1, Polysciences) on top of the media in each well, enough for each 6 well column with a 10% extra margin, as described previously(*72*). Cells were left to incubate overnight, for a minimum of 12 hours. The next morning media was replaced with fresh media including the respective dilutions of the inducing agent.

Cellular detection assay

For secreted NanoLuc assays, cells were plated on clear 96 well cell culture plates (655-180, Greiner Bio-One). The next day cells were transfected with STAT3 (100ng), STAT3-NanoLuc reporter (150ng), and either a single GEMS receptor chain containing Bcl2 or DBVen 1619_V2 (600ng) or both chains together (300ng each). The following day cells were induced with different dilutions of the inducing agent Venetoclax. After 24 hours of induction, 5uL media was transferred to a black 384 well plate (3820, Corning) and mixed with 5uL diluted substrate from the Nano-Glo Luciferase Assay kit (N1120, Promega). After gentle shaking, plates were measured on a Tecan Spark plate reader with an integration time of 1000ms.

For intracellular NanoLuc assays, cells were plated in black 96 well cell culture plates (655086, Greiner). The next day cells were transfected with either a single chain of PDF1-C-term-NanoLuc or DBAct553_1-N-term-NanoLuc (825ng) or both chains together (412.5ng each). The following day cells were induced with different dilutions of the inducing agent Actinonin. After 24 hours of induction, intracellular Nanoluciferase activity was measured using the Nano-Glo Live Cell Assay kit (N2012, Promega). Media was aspirated and replaced with 24uL RPMI Medium (52400-025, Gibco) containing 10% v/v FBS and 6uL diluted substrate was added to each well. After gentle shaking, plates were measured on a Tecan Spark plate reader with an integration time of 1000 ms. All cell-based fits presented in Fig. 5 were calculated from technical replicates (n = 3) using a nonlinear four-parameter curve fitting analysis. All statistical analyses are based on a two way ANOVA with multiple comparisons. Data points represent technical replicates (n=3) with mean and standard deviation.

Cell-free reporter system

The gene encoding the 6xHis-DBPro1156_2 protein fused to T7 RNA Polymerase (T7RNAP) was cloned into a pQE30 plasmid using Gibson assembly. The plasmid was then transformed into NEBExpress Iq competent *E. coli* (NEB; ref: C3037I) for protein expression. Bacteria were pre-cultured overnight and inoculated to a 500 ml LB-medium culture, grown until the $OD_{600}$~0.7, and then induced with 0.1 mM IPTG for 3 hours. The cells were collected by centrifugation at 4000 g and lysed by sonication. Proteins were purified using Ni-NTA IMAC sepharose gravity columns.

The ZF438-DB3 scFv ($V_H$/$V_L$) fusion protein was expressed using a PURExpress kit from NEB (E6800S) with the addition of a disulfide bond enhancer (E6820S). The reaction volume was 10 μl, containing 4 μl of solution A, 3 μl of solution B, 0.4 μl of NEB disulfide bond enhancer 1, 0.4 μl of NEB disulfide bond enhancer 2, 2 μl of DNA template (10 ng/μl), and 0.2 μl of water. The reaction was incubated at 34 ˚C for 3 hours and used for the following reporter reaction.

PURExpress kit from NEB (E6800S) with disulfide bond enhancer (E6820S) was used to set up the mCherry reporter expression as well. The reporter-expressing reaction additionally includes 100 nM purified DBPro1156_2-T7RNAP and ZF438-DB3 scFv pre-expressed with PURExpress. DNA template for the mCherry gene is set to 4 nM, the mCherry gene is transcribed under the regulation of a truncated T7 promoter downstream of the zinc finger 438 protein binding site, which requires a zinc finger protein for activating transcription. Progesterone was dissolved in 2% DMSO. 10 μl reactions with different conditions are loaded into a 384-well plate. The mCherry fluorescent intensity is measured on a BioTek Synergy H1 Multimode Reader (Agilent) with an excitation wavelength of 565 nm and an emission wavelength of 615 nm at 34 ˚C for 8 hours with 2-minute intervals. All cell-based fits presented in Fig. 5 were calculated from technical replicates (n = 3) using a nonlinear four-parameter curve fitting analysis. All statistical analyses are based on a two way ANOVA with multiple comparisons. Data points represent technical replicates (n=3) with mean and standard deviation.

Protein purification for crystallography

6xHis-tagged PDF1 from Pseudomonas aeruginosa and DBAct553_1 were expressed in E. coli (BL21 T7 Express). Amino acid sequences of both proteins are shown in table S5. For PDF1, cells were grown in Luria-Bertani (LB) medium supplemented with 100mM $NiSO_4$ up to an $OD_{600}$ of 0.7 at 37°C, induced with 1mM IPTG and continued growing overnight at 18°C. For DBAct553_2, cells were grown in auto-induction medium (AIM) up to $OD_{600}$ of 0.7 at 37°C and then overnight at 18°C. Cells were collected by centrifugation at 4000g for 10 min, resuspended in lysis buffer (50 mM Tris, pH 7.5, 500 mM NaCl, 5% glycerol, 1 mg $ml^{-1}$ lysozyme, 1 mM PMSF and 1 μg $ml^{-1}$ DNase) and lysed by sonication. Lysates were then clarified by centrifugation at 30'000g for 30 min and filtered. Proteins were purified using the ÄKTA pure system (GE healthcare) Ni-NTA HisTrap affinity column followed by a size exclusion chromatography on a Superdex HiLoad 16/600 75pg with TBS (50mM Tris pH 7.5, 250mM NaCl, 10μM $NiSO_4$) as a final buffer. PDF1, DBAct553_2 and Actinonin were mixed at a final concentration of 35µM, 105µM and 300µM respectively and incubated on ice for 1 hour. Proteins were then concentrated by centrifugation prior to crystallization.

Crystallographic data collection and structure determination

The Actinonin-bound PDF1:DBAct553_1 complex (5 mg/ml) was crystallized using the sitting drop vapor diffusion setup at 18°C with 200nl of protein and 200nl crystallization solution consisting of 0.2M sodium formate, 0.1M sodium phosphate pH 6.2, 20% (v/v) PEG and 10% (v/v) glycerol. Crystals were cryoprotected with 25% glycerol and flash-cooled in liquid nitrogen. Diffraction data were collected at a temperature of 100K at the European Synchrotron Radiation Facility (ESRF Grenoble, France). Raw data were processed and scaled with XDS, and then processed using the autoPROC package(*73*). Phases were obtained by molecular replacement using the Phaser module of the Phenix package and a model from PDB 1LRY in complex with our designed binder DBAct553_1(*74*). Atomic model adjustment and refinement was completed using COOT and Phenix.refine(*75*, *76*). Finally, MolProbity(*77*) was used to assess the quality of the refined model. Details of data collection and refinement statistics are shown in table S6.

Cryo-EM preparation and data acquisition

A chimeric DB3 Fab (see table S5) was produced using the Expi293 expression system from Thermo Fisher Scientific (A14635). An anti-kappa light chain Fab(*78*) (see table S5) was produced using the ExpiCHO-S cells (Thermo Fisher Scientific, ref: A29127) growing in a ProCHO-5 medium (Lonza) supplemented with 2% DMSO. Supernatants were collected 6 and 7 days respectively after transfection, filtered and purified by Ni-NTA affinity chromatography followed by a size exclusion chromatography on a Superdex HiLoad 16/600 75pg. All proteins were concentrated in PBS as a final buffer. DBPro1156_2 was purified as indicated previously in the “protein expression and purification” section.

DB3 Fab, anti-kappa light chain Fab, DBPro1156_2 and progesterone were mixed with a molar ratio of 1 : 0.9 : 3 : 2 respectively, supplemented with 0.1% n-dodecyl-ß-D-maltoside (DDM) and concentrated to 3.87 mg/ml. Proteins were applied to a glow discharged 300-mesh holey carbon grid (Au 1.2/1.3 Quantifoil Micro Tools), blotted for 4 s at 95% humidity, 10 °C, plunge frozen in liquid ethane (Vitrobot, Thermo Fisher Scientific(TFS)) and stored in liquid nitrogen. Data collection was performed on a 200 kV Glacios cryo-EM microscope using EPU 3.7.0 (TFS) equipped with a Falcon4i detector. Micrographs were recorded at a calibrated magnification of 150000 with a pixel size of 0.926 Å and a nominal defocus ranging from -1.2 µm to -2.4 µm.

Cryo-EM image processing

Acquired cryo-EM data was processed using cryoSPARC v4.4.1. Particles were picked from motion and gain corrected micrographs using a blob picker and extracted with a box size of 360 pixels, downsampled to 180. Following 2D classification and *ab initio* 3D reconstruction, clean particles were used for non-uniform 3D reconstruction to obtain a low-resolution map of ~6 Å. For structure building, the *in silico* models generated in our study were split into segments and docked into density using ChimeraX(*79*) and rigid body refined using Coot(*75*).

Chemical synthesis

All chemical reagents and solvents for synthesis were purchased from commercial suppliers (Sigma-Aldrich, Fluka, Acros) and were used without further purification or distillation. The composition of mixed solvents is given by the volume ratio (v/v). $^{1}$H nuclear magnetic resonance (NMR) spectra were recorded on a Bruker DPX 400 (400 MHz for $^{1}$H) with chemical shifts (δ) reported in ppm relative to the solvent residual signals (7.26 ppm for of $CDCl_3$; 3.31 ppm for MeOD) (fig. S14). Coupling constants are reported in Hz. LC-MS was performed on a Shimadzu MS2020 connected to a Nexerra UHPLC system equipped with a Waters ACQUITY UPLC BEH Phenyl 1.7µm 2.1x50mm column. Buffer A: 0.05% HCOOH in $H_2O$ Buffer B: 0.05% HCOOH in acetonitrile. LC gradient: 10% to 90% B within 6.0 min with 0.5 ml/min flow. Preparative HPLC was performed on a Dionex system equipped with an UltiMate 3000 diode array detector for product visualization on a Waters SymmetryPrep C18 column (7 µm, 7.8 x 300 mm). Buffer A: 0.1% v/v TFA in $H_2O$; Buffer B: acetonitrile. Gradient was from 25% to 90% B within 30 min with 3 ml/min flow.

*19-O-benzoylprogesterone*

19-hydroxyprogesterone (2.0 mg, 6.1 µmol, 1 eq.) was dissolved in pyridine (0.5 ml) and benzoyl chloride (0.9 ul, 7.9 µmol, 1.3 eq) was added. The reaction mixture was stirred for 3h. LC-MS analysis showed reaction completion and 10 µl methanol were added. After 30 minutes, the solvents were evaporated under reduced pressure. The residue was dissolved in a minimum of acetonitrile and subjected to preparative HPLC. The fractions containing the product were pooled and lyophilized. Yield: 1.1 mg (41%). $^{1}$H NMR (400 MHz, $CDCl_3$) δ 7.89 (d, J = 8.4 Hz, 2H), 7.56 (t, J = 7.4 Hz, 1H), 7.42 (t, J = 7.8 Hz, 2H), 5.98 (s, 1H), 4.81 (d, J = 11.3 Hz, 1H), 4.46 (d, J = 11.3 Hz, 1H), 2.68 (ddd, J = 17.0, 13.8, 5.9 Hz, 1H), 2.57 – 2.32 (m, 4H), 2.26 – 2.06 (m, 5H), 2.03 – 1.63 (m, 6H), 1.55 – 1.37 (m, 2H), 1.36 – 1.06 (m, 5H), 0.69 (s, 3H). HRMS (ESI/QTOF) m/z: $[M+H]^+$ Calcd for $C_{28}H_{35}O_4^+$ 435.2530; Found 435.2528.

*TBDMS-Actinonin*

Actinonin (2.0 mg, 5.2 µmol, 1 eq.) and 4-dimethylaminopyridine (3.8 mg, 31.2 µmol, 6 eq.) were suspended in DCM (0.5 ml). TBDMS-Cl (2.5 mg, 16.6 umol, 3.2 eq.) was added and the reaction was stirred for 5h at r.t. The solvent was evaporated under reduced pressure, the residue was dissolved in MeOH (0.5 ml), water (50 µl) was added and the reaction was heated to 60°C for 5h. The solvents were evaporated again, the residue was dissolved in a minimum of DCM and subjected to preparative TLC using DCM/MeOH 9:1 as the eluent. Yield: 2.0 mg (77%). 1H NMR (400 MHz, MeOD) δ 4.38 (d, J = 8.5 Hz, 1H), 4.13 (s, 1H), 3.89 (dt, J = 10.0, 6.8 Hz, 1H), 3.79 (dd, J = 9.9, 5.3 Hz, 1H), 3.68 (dd, J = 9.9, 2.8 Hz, 1H), 3.63 – 3.42 (m, 1H), 2.83 – 2.75 (m, 1H), 2.34 (dd, J = 14.5, 8.0 Hz, 1H), 2.24 – 1.84 (m, 6H), 1.67 – 1.48 (m, 1H), 1.46 – 1.18 (m, 6H), 1.02 – 0.94 (m, 7H), 0.93 – 0.86 (m, 12H), 0.07 (s, 3H), 0.05 (s, 3H). HRMS (ESI/QTOF) m/z: $[M+Na]^+$ Calcd for $C_{25}H_{49}N_3NaO_5Si^+$ 522.3334; Found 522.3342.

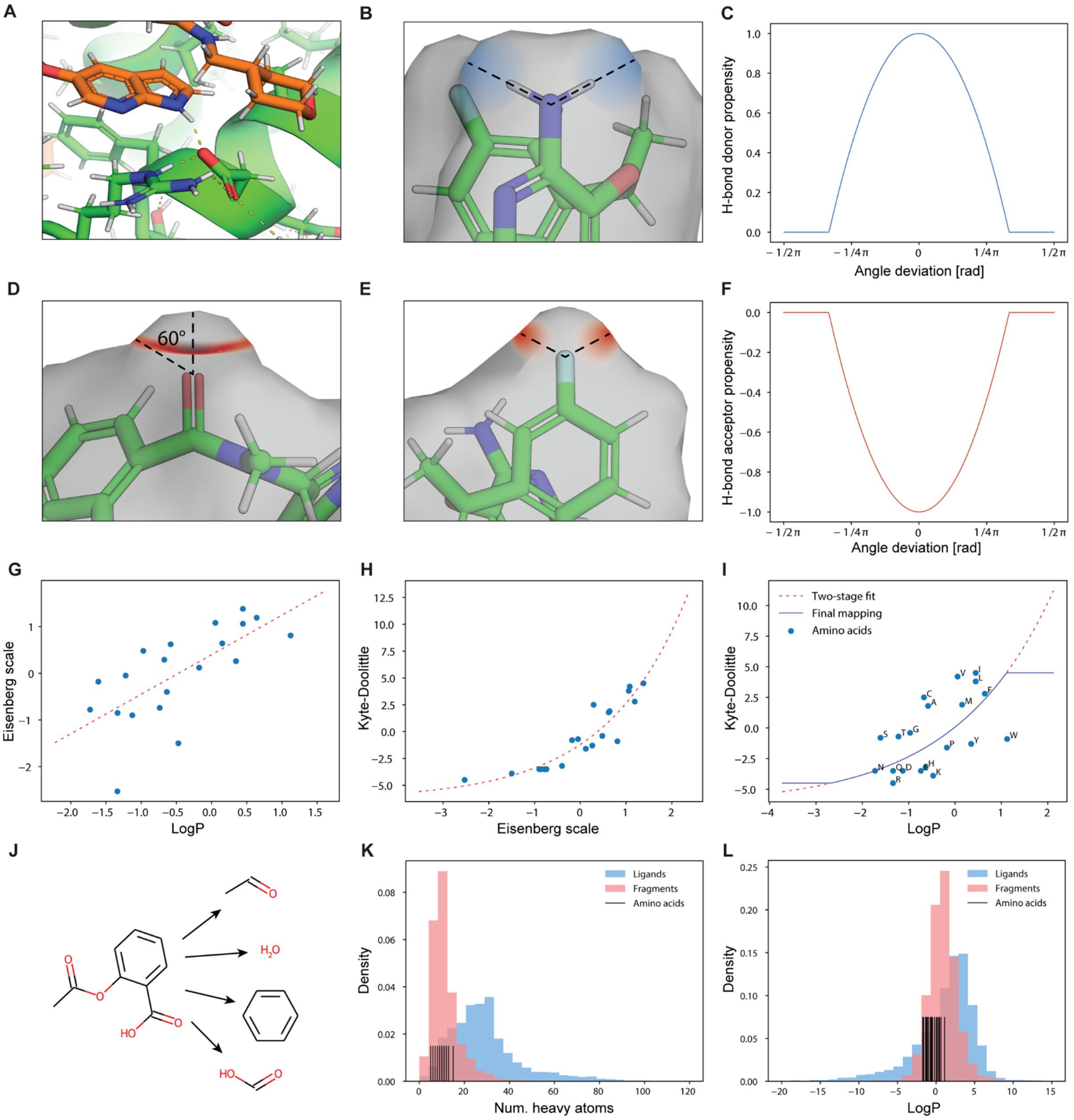


**Fig. S1: MaSIF feature computation for small molecule ligands. A-F.** Hydrogen bond propensity is assigned in a direction-dependent manner(*34*). Surface points are assigned positive (donor) or negative (acceptor) values based on their distance to the ideal direction (C, F). The optimal position for an acceptor can either lie anywhere on a cone (D) or in one or more unique directions (E). **G-I.** For hydropathy, we convert computational LogP values to the protein-specific Kyte-Doolittle (KD) scale required by MaSIF using the Eisenberg scale as an intermediate. We also restrict the outputs to be between the minimum and maximum KD values after the mapping leading to the relationship shown in panel (I). **J-L.** Ligands were fragmented into smaller objects (J). The reason is that most ligands are significantly larger than any amino acid (K) and exhibit more extreme LogP values (L). We therefore compute hydrophobicity values based on fragments. This procedure ensures that the new hydropathy feature remains “in-distribution” of the pre-trained MaSIF model.

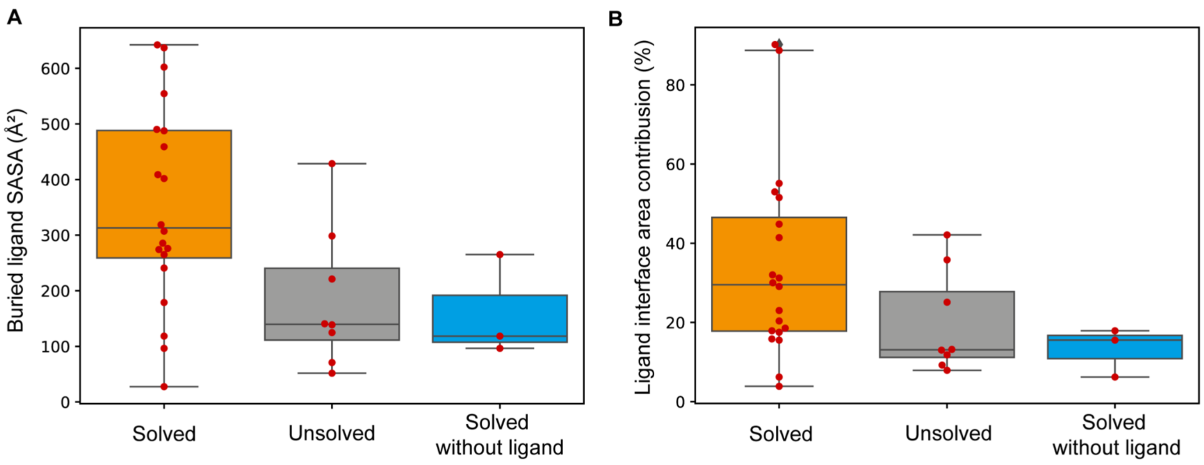


**Fig. S2: Ligand interface area contribution in the benchmark dataset. A-B.** Buried solvent-accessible surface area (SASA) contribution of the ligand in absolute value (A) or in percentage of the total interface surface area (B) for the 28 protein-ligand complexes used in the benchmark dataset of known ternary complexes. Complexes were categorized based on the benchmark outcome, namely successfully solved complexes with the ligand (orange), unsolved complexes with the ligand (gray) or solved without the ligand (blue).

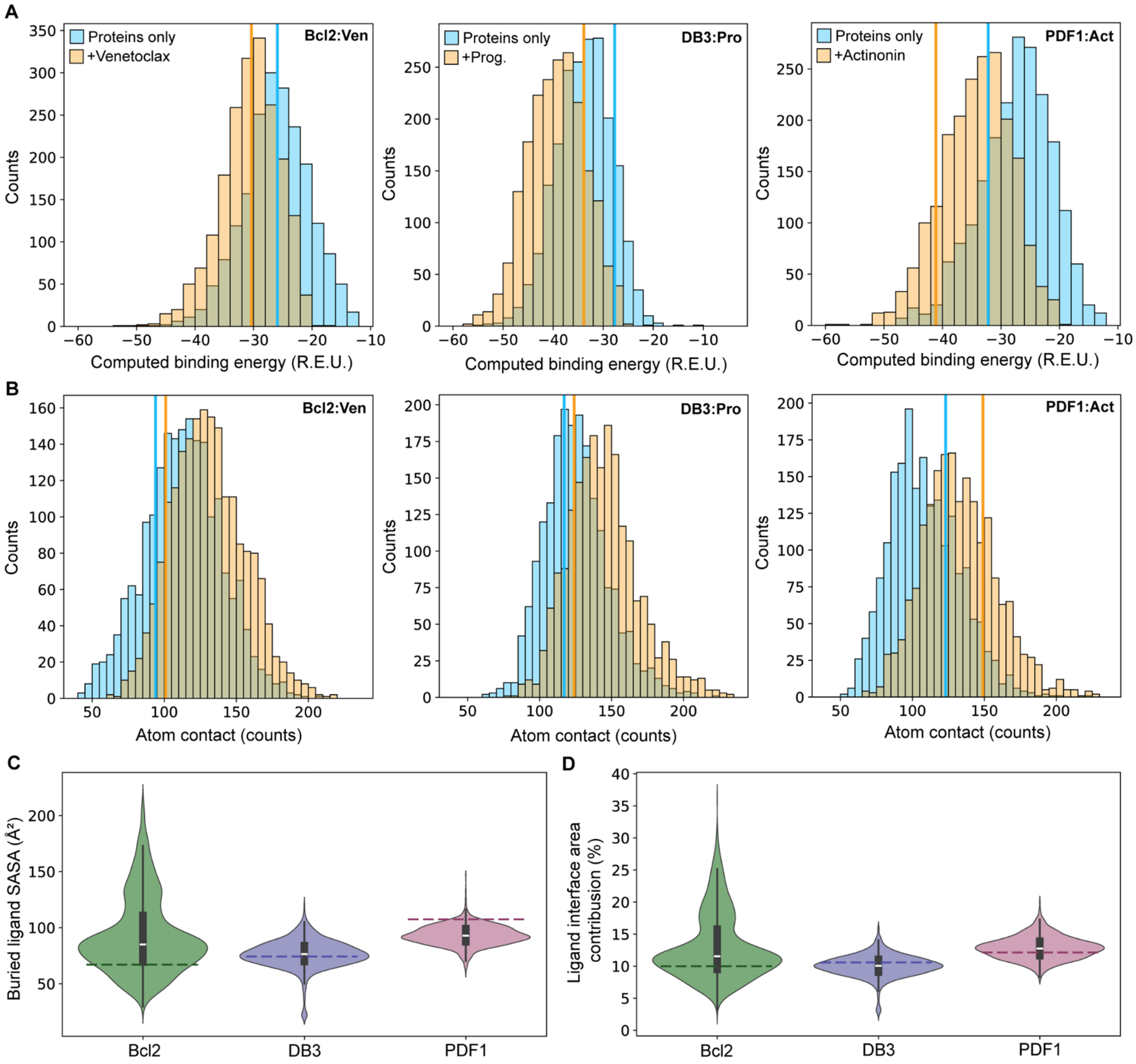


**Fig. S3: Neosurface properties captured by the designed binders. A-B.** Computed binding energy (ΔΔG, A) and number of atomic contacts (B) for all designs targeting Bcl2:Venetoclax (left), DB3:Progesterone (middle) and PDF1:Actinonin (right) complexes. Calculations were done in absence (blue) and presence (orange) of the respective small molecules. Atom contacts were defined based on the Van der Waals radii ($r_{VdW}$ + 0.2Å tolerance) of each pair of atoms. Vertical lines represent the identified binder for each targeted complex. **C-D**. Buried solvent-accessible surface area (SASA) of the ligand (C) and ligand contribution with respect to the total buried SASA (D) for each target protein-ligand complex. Dashed lines represent the identified binder for each targeted complex.

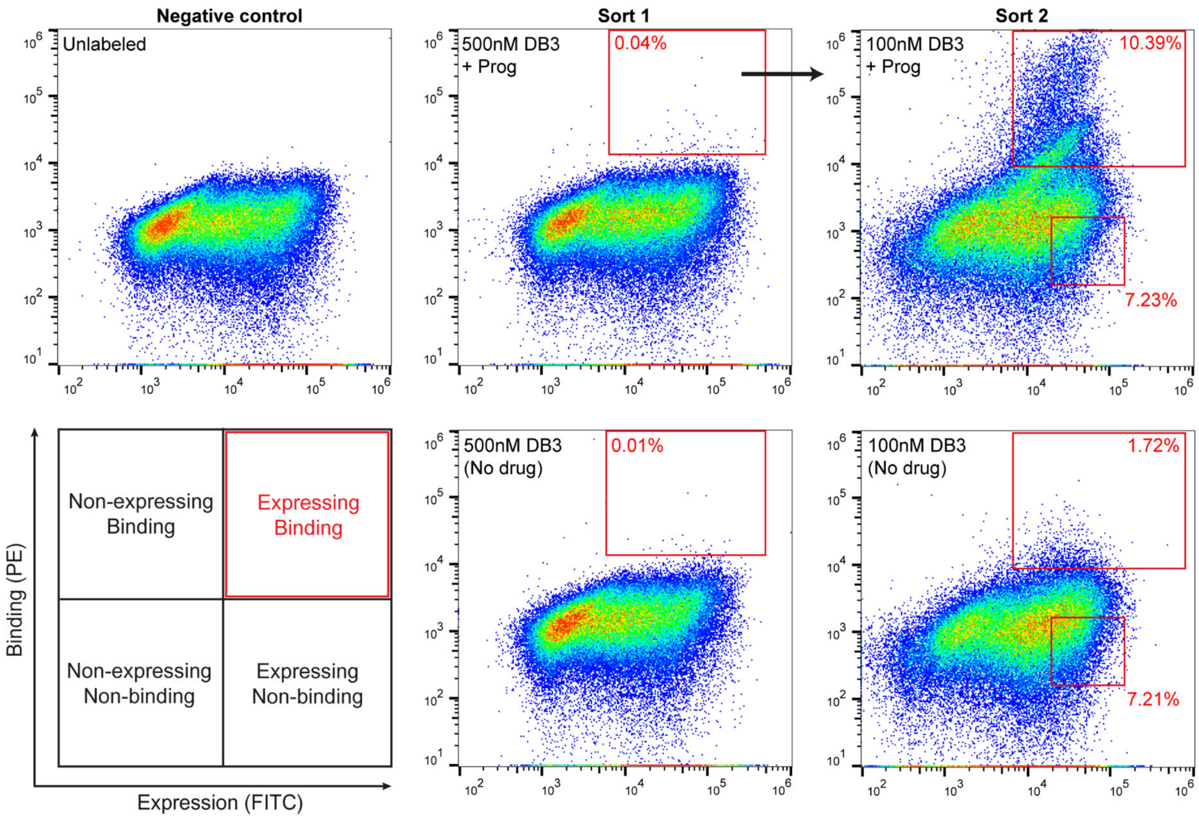


**Fig. S4: Representative flow cytometry graphs of the binder screening.** Yeast surface display screening of the first and second sort of the library against DB3:Progesterone in presence (+Prog) and absence of the small molecule. Yeasts labeled with secondary antibodies but without any ligand were used as a negative control to set the gates. In sort 1, binding population from the selected gates was used for a second sort. In sort 2, yeasts were sorted for both binding (upper gate) and non-binding (lower gate) populations and used for next-generation sequencing.

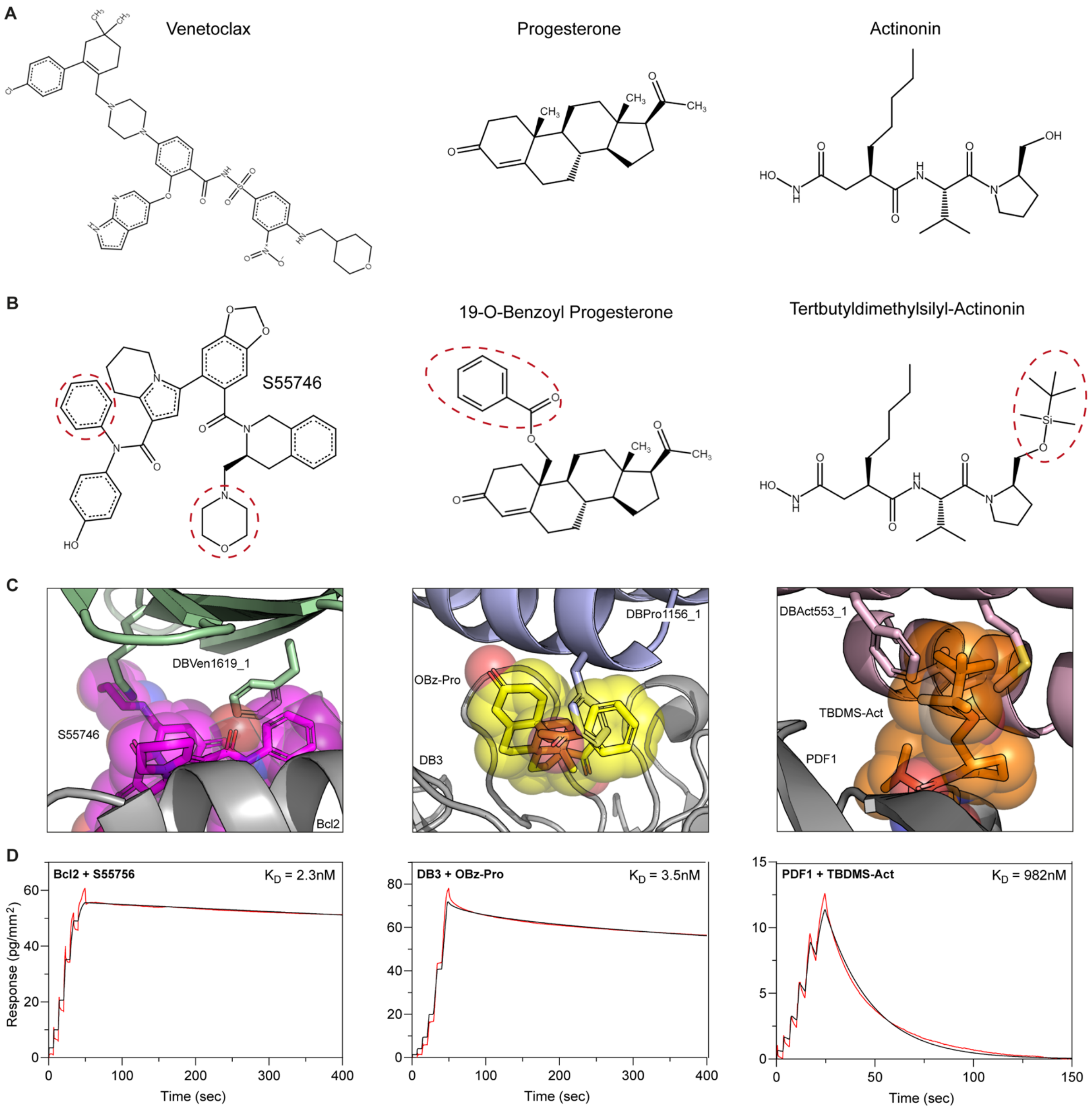


**Fig. S5: Binding control of small molecules analogs. A.** Original small molecule used for each drug:protein complex. **B.** Small molecule derivative binding to the same target but introducing a clash with the designed binder. Steric clashes are indicated with red dashed circles. **C.** Crystal structure (S55746; PDB: 6GL8) or computational models (19-O-Benzoyl Progesterone and Tertbutyldimethylsilyl-Actinonin) of the small molecule analogs relaxed by Gnina(*80*) with their respective target protein in absence of the designed binders. Complexes were then overlapped with the computational model of the designed binders to identify clashing regions. **D.** WaveRAPID binding kinetics of each small molecule analog to their respective target protein performed by Grating-Coupled Interferometry (GCI). Measurements are indicated in red and fit curves in black..

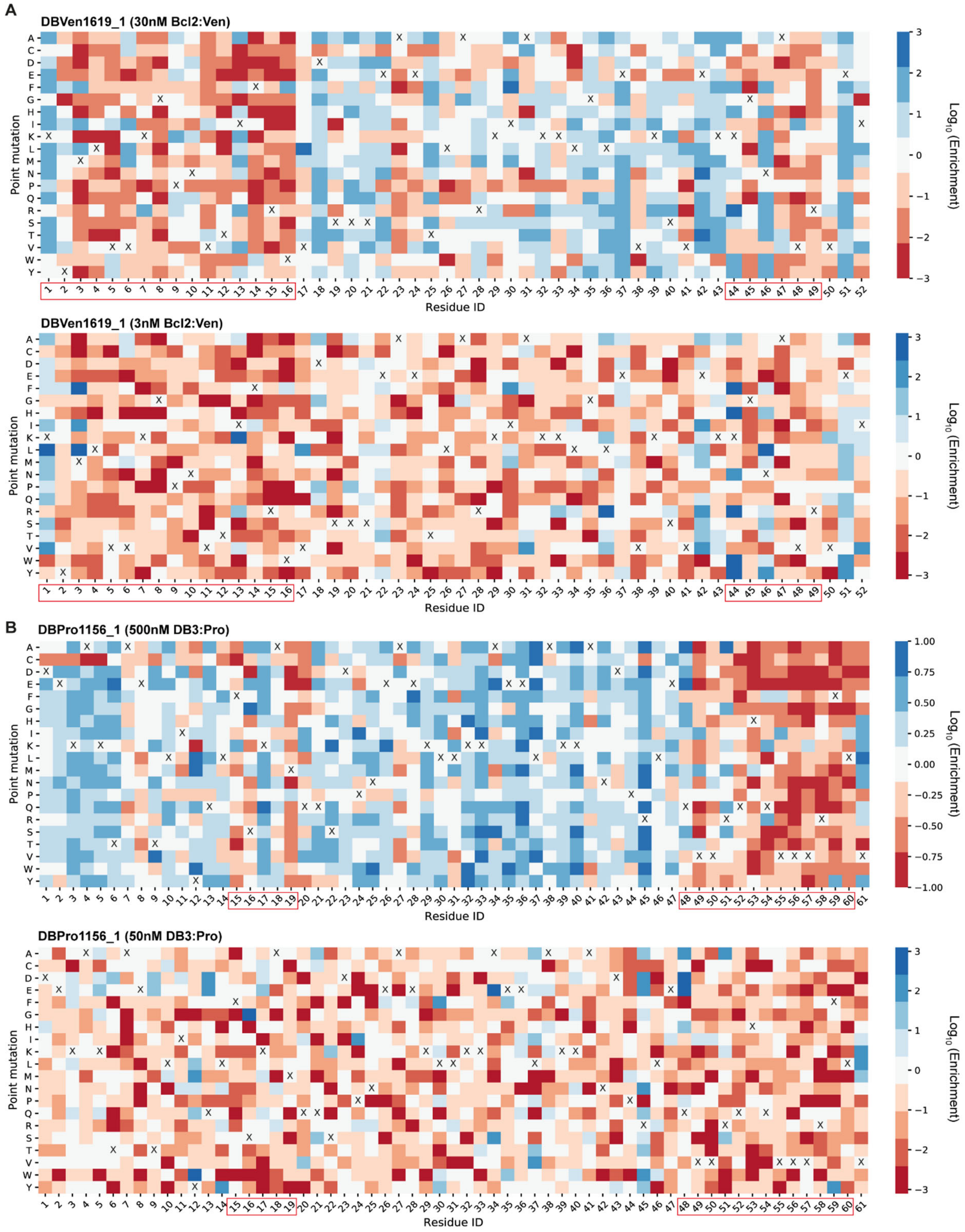

A
DBVen1619_1 (30nM Bcl2:Ven)
DBVen1619_1 (3nM Bcl2:Ven)
B
DBPro1156_1 (500nM DB3:Pro)
DBPro1156_1 (50nM DB3:Pro)
Point mutation
Residue ID
Log10 (Enrichment)

C

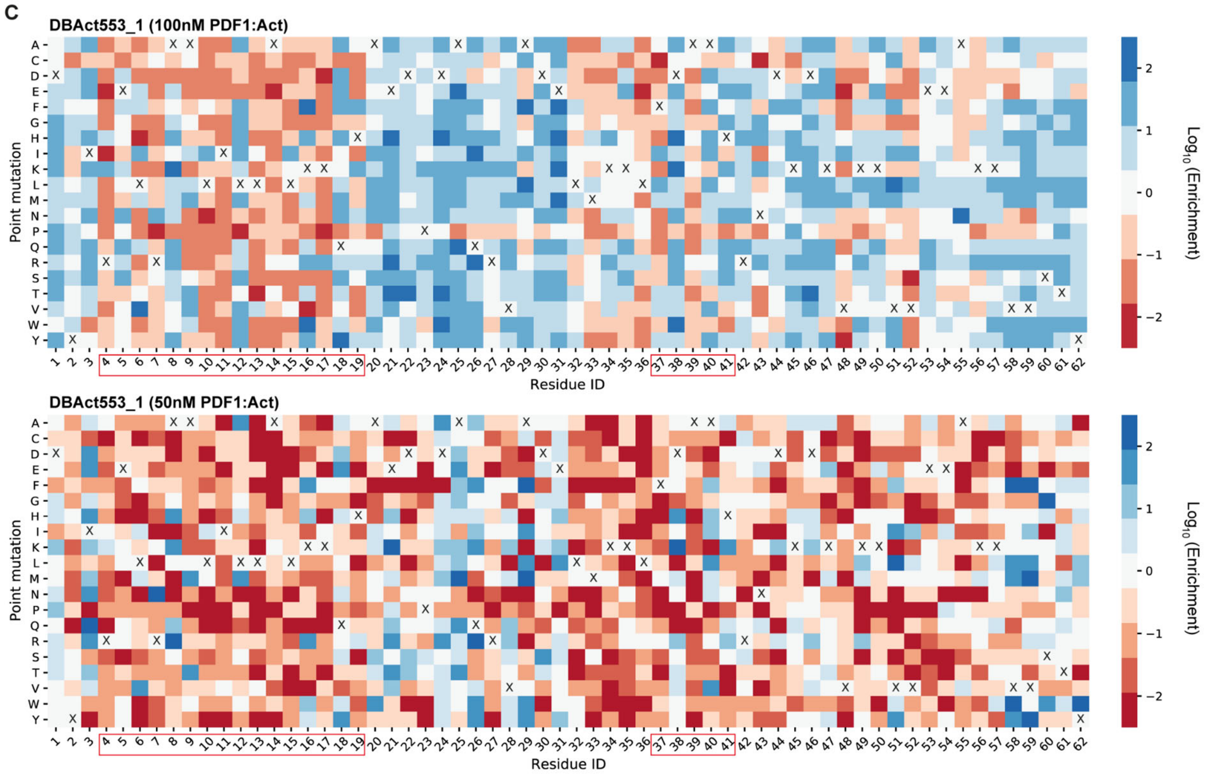


**Fig. S6: Full data of the site-saturation mutagenesis. A-C.** Heatmaps represent the logarithmic value of the enrichment score (Counts in binding population divided by counts in non-binding population) of each mutation at every position. Sorting has been performed following the gating strategy presented in fig. S2. Native amino acids are marked with a cross (X) and near-interface positions are highlighted with a red box. Sortings were performed with a high concentration and a low concentration of target complex to focus on deleterious and beneficial mutations respectively. Site saturation mutagenesis were performed on DBVen1619_1 for the the Bcl2:Venetoclax complex (A), DBPro1156_1 for the DB3:Progesterone complex (B) and DBAct553_1 for the PDF1:Actinonin complex (C).

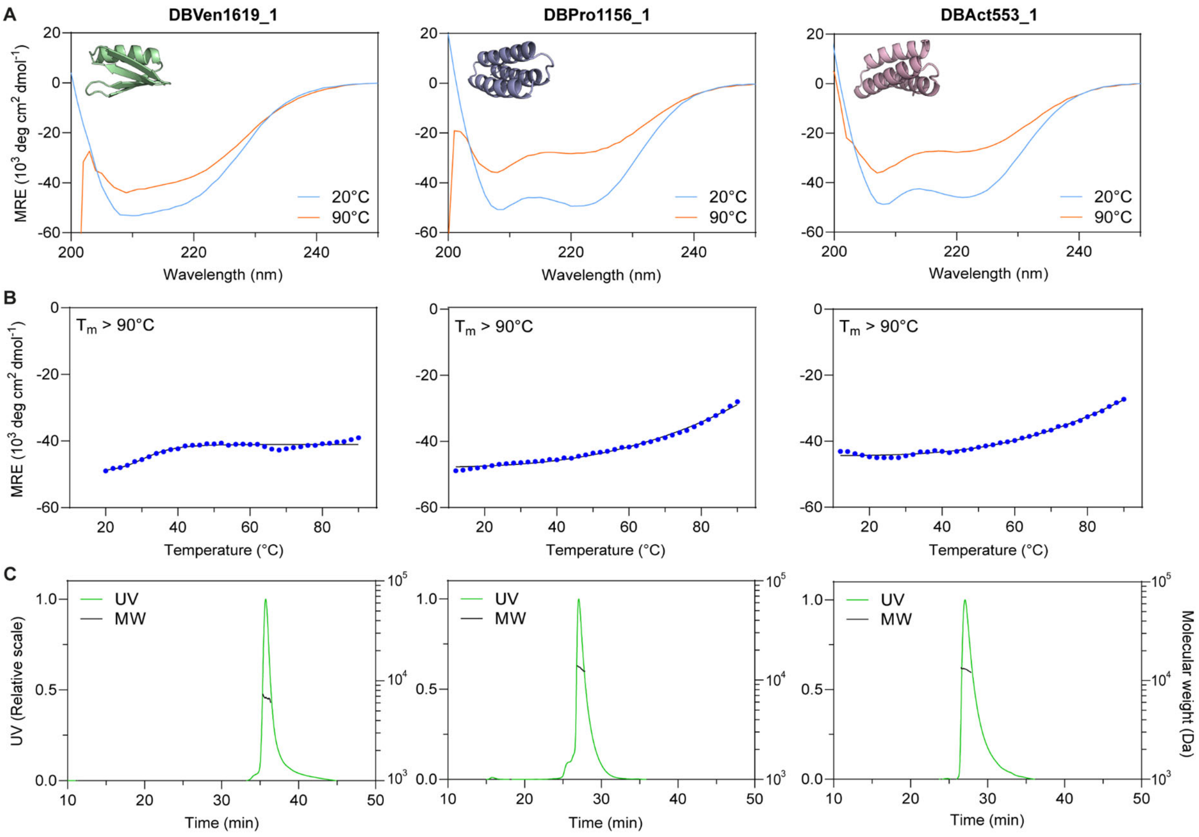


**Fig. S7: Biophysical characterization of purified binders. A.** Protein folding of the purified binder measured by circular dichroism at 20°C (blue) or 90°C (orange). **B**. Thermal stability determined by measuring the ellipticity at 218nm at increasing temperature. **C.** Oligomeric state determined by size-exclusion multi-angle light scattering (SEC-MALS)

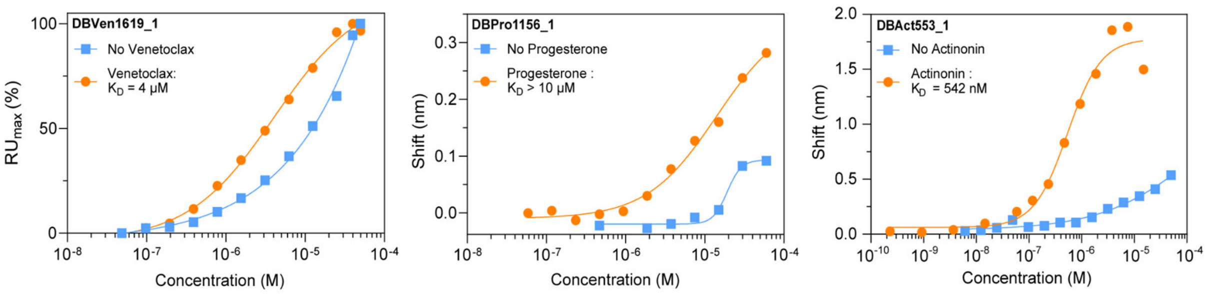


**Fig. S8: Affinity measurements of first-generation purified binders.** Affinity measurement for DBVen1619_1, DBPro1156_1 and DBAct553_1 performed by surface plasmon resonance (DBVen1619_1) or biolayer interferometry (DBPro1156_1 and DBAct1156_1). Each measurement was performed in presence (orange) or absence (blue) of the respective small molecule. The fits were calculated using a nonlinear four-parameter curve fitting analysis.

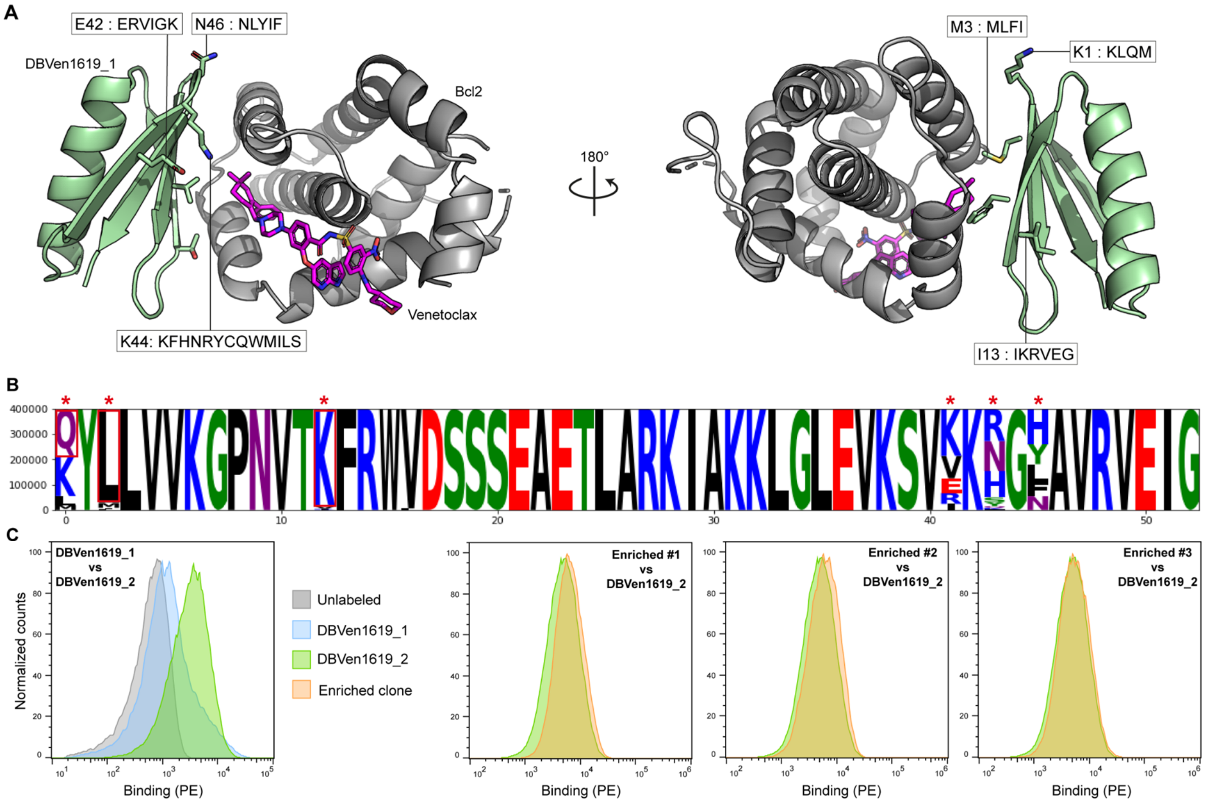


**Fig. S9: Experimental optimization of DBVen1619. A.** Computational model of DBVen1619_1 (green) in complex with Bcl2 (gray) and Venetoclax (Magenta). Potential beneficial mutations obtained from site-saturation mutagenesis (SSM) data and subsequent degenerate codons are found in black boxes for each mutated position. **B.** Sequence logo plot of the combinatorial library sorted twice with yeast display. Mutated positions are highlighted with a red asterisk. Mutations selected to constitute DBVen1619_2 are highlighted with a red square. **C.** Comparison of unlabeled yeast (gray), or yeasts displaying DBVen1619_1 (blue), DBVen1619_2 (green, K1Q+M3L+I13K) or the top 3 most enriched sequences of the combinatorial library (orange). All yeasts were labeled with 3nM Bcl2:Venetoclax complex.

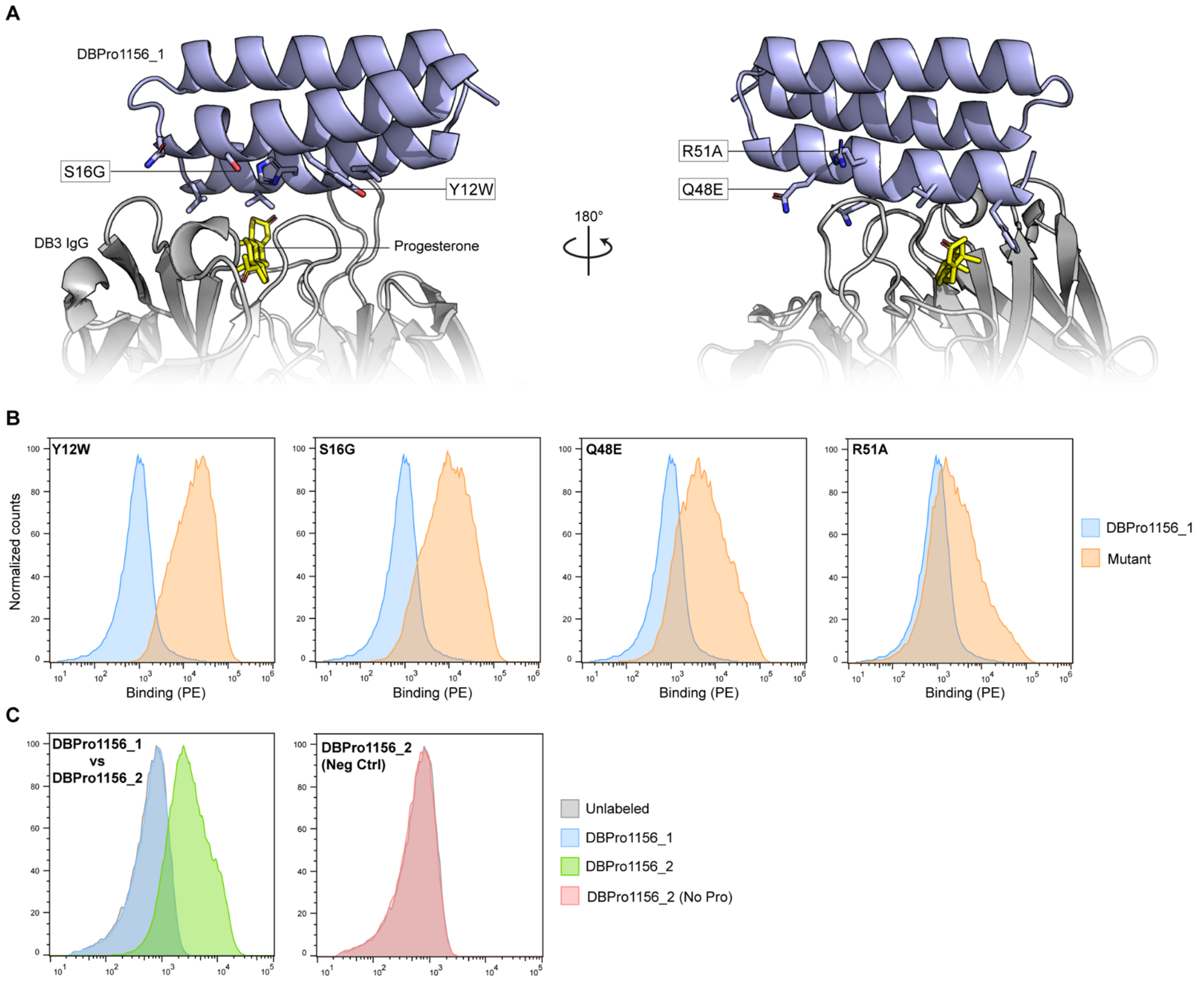


**Fig. S10: Experimental optimization of DBPro1156. A.** Computational model of DBPro1156_1 (blue) in complex with DB3 IgG (gray) and Progesterone (yellow). Potential beneficial mutations obtained from site-saturation mutagenesis (SSM) data are found in black boxes for each mutated position**. B.** Binding signal measured by flow cytometry for yeasts displaying DBPro1156_1 (blue) or the corresponding mutant (orange). All yeasts were labeled with 50nM DB3:Progesterone complex. **C.** Binding signal measured by flow cytometry measured for yeasts displaying DBPro1156_1 (blue) compared to DBPro1156_2 (green, Y12W+S16G). Yeasts were labeled with 3nM DB3:Progesterone complex. Negative control performed on yeast displaying DBPro1156_2 labeled with DB3 only.

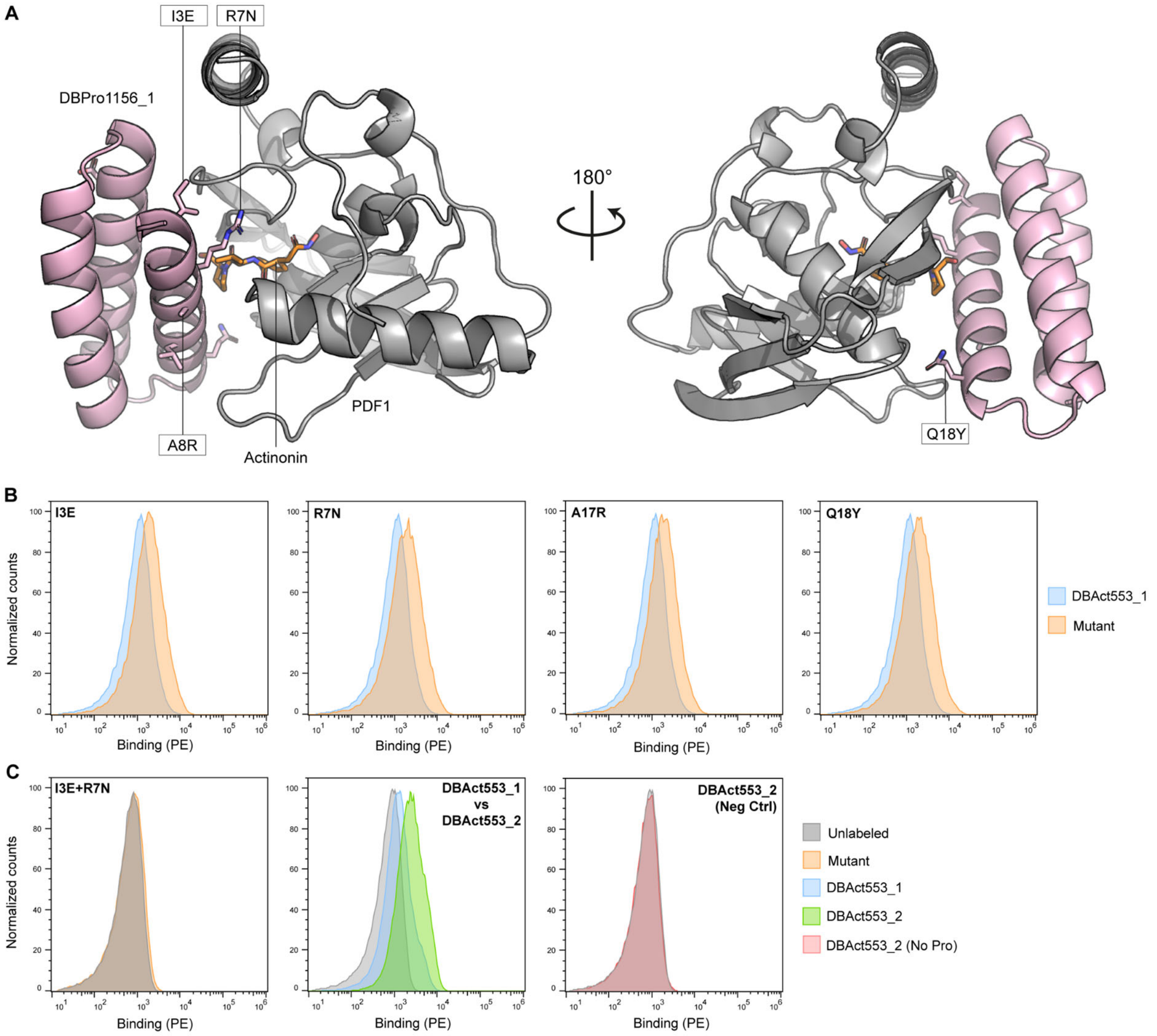


**Fig. S11: Experimental optimization of DBAct553. A.** Computational model of DBAct553_1 (pink) in complex with PDF1 (gray) and Actinonin (orange). Potential beneficial mutations obtained from site-saturation mutagenesis (SSM) data are found in black boxes for each mutated position**. B.** Binding signal measured by flow cytometry for yeasts displaying DBAct553_1 (blue) or the corresponding mutant (orange). All yeasts were labeled with 10nM PDF1:Actinonin complex. **C.** Binding signal measured by flow cytometry measured for yeasts displaying DBAct553_1 (blue) compared to DBAct553_2 (green, R7N+A8R). Yeasts were labeled with 10nM PDF1:Actinonin complex. Negative control performed on yeast displaying DBAct553_2 labeled with DB3 only. Combination of mutants I3E+R7N was deleterious for binding as compared to unlabeled yeasts.

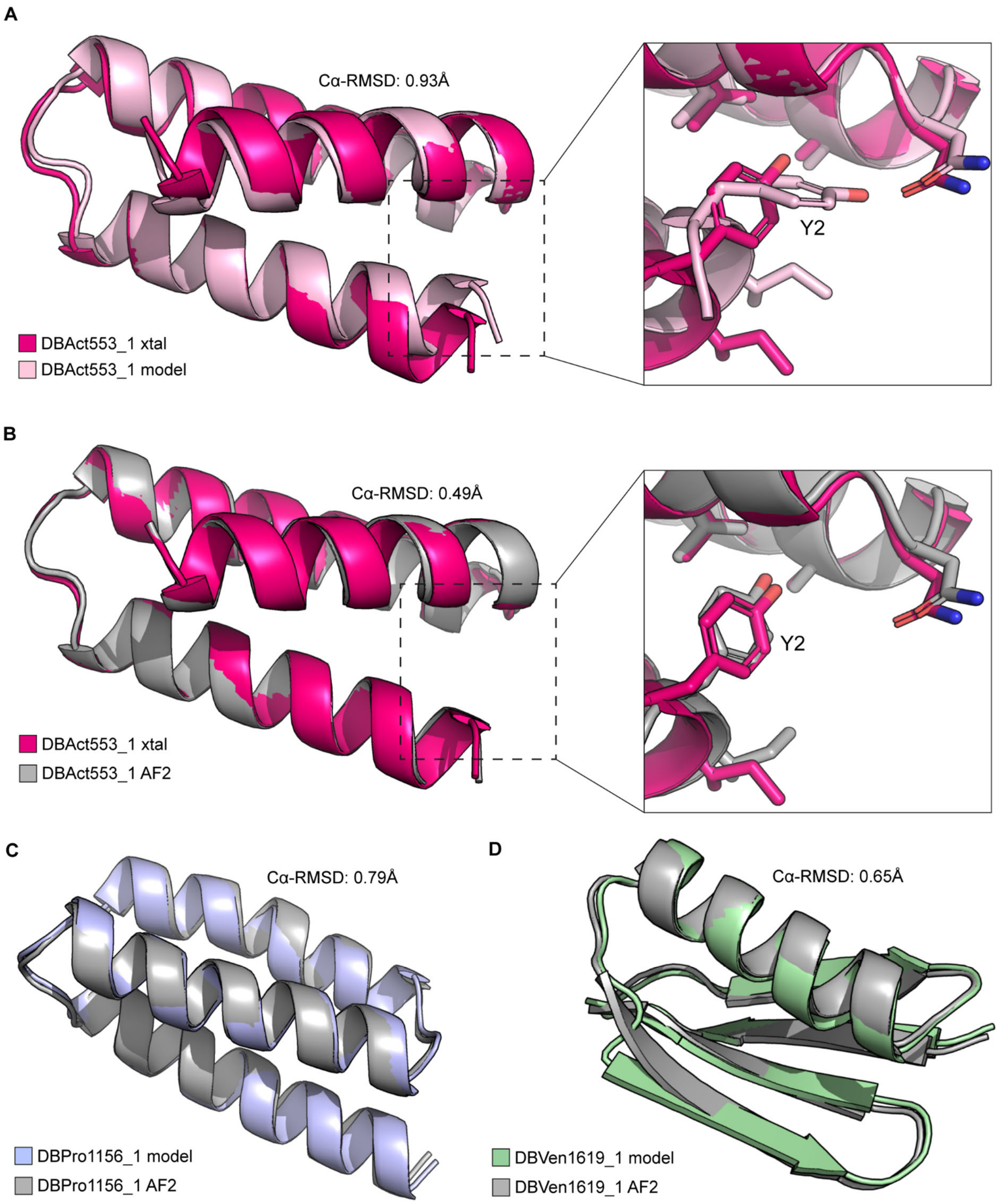


**Fig. S12: Comparison between crystallographic data and AlphaFold2 predictions. A.** Computational model of DBAct553_1 (light pink) aligned with its crystal structure (magenta) with a close-up on tyrosine-2. **B.** AlphaFold2 (AF2) prediction of DBAct553_1 (gray) aligned with its crystal structure (magenta) with a close-up on tyrosine-2. **C-D**. Comparison between computational models of DBPro1156_1 (C) and DBVen1619_1 (D) and their respective AlphaFold2 prediction as monomers.

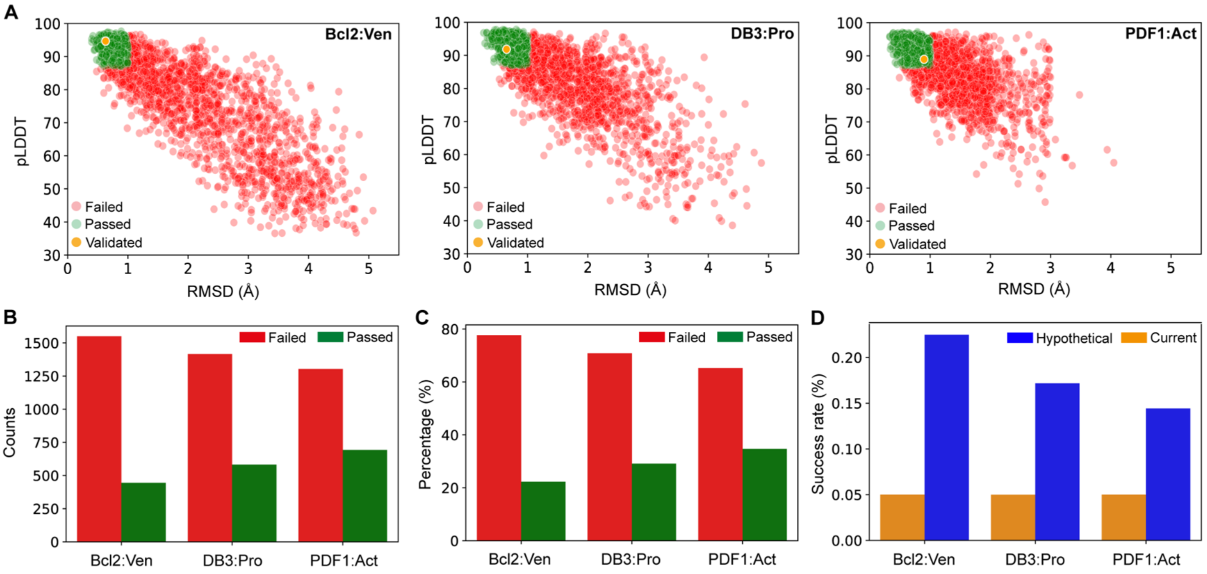


**Fig. S13: AlphaFold prediction and post-filtering of generated designs. A.** AlphaFold monomer prediction (single sequence mode) of the ~2000 designs generated against each drug:protein complex. Prediction confidence (pLDDT) and root mean square deviation (RMSD) from the computational models are plotted. Designs that would pass a strict filtering (RMSD ≤ 1Å and pLDDT ≥ 87) are colored in green, while ones that failed filtering are colored in red. Validated binders are colored in orange. **B-C.** Counts (B) and percentage (C) of generated designs that failed (red) or passed (green) the strict AlphaFold2 filtering. **D.** Experimental success rate obtained with the current data (orange) compared to the hypothetical success rate (blue) if a strict filtering with AlphaFold2 (RMSD ≤ 1Å and pLDDT ≥ 87) was used prior screening.

A

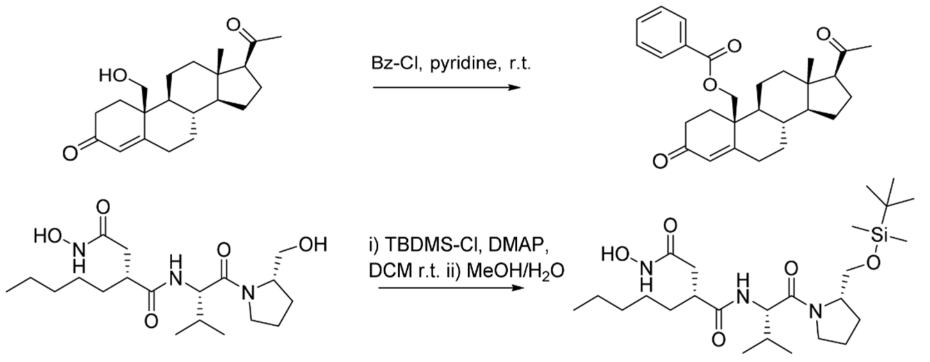


B

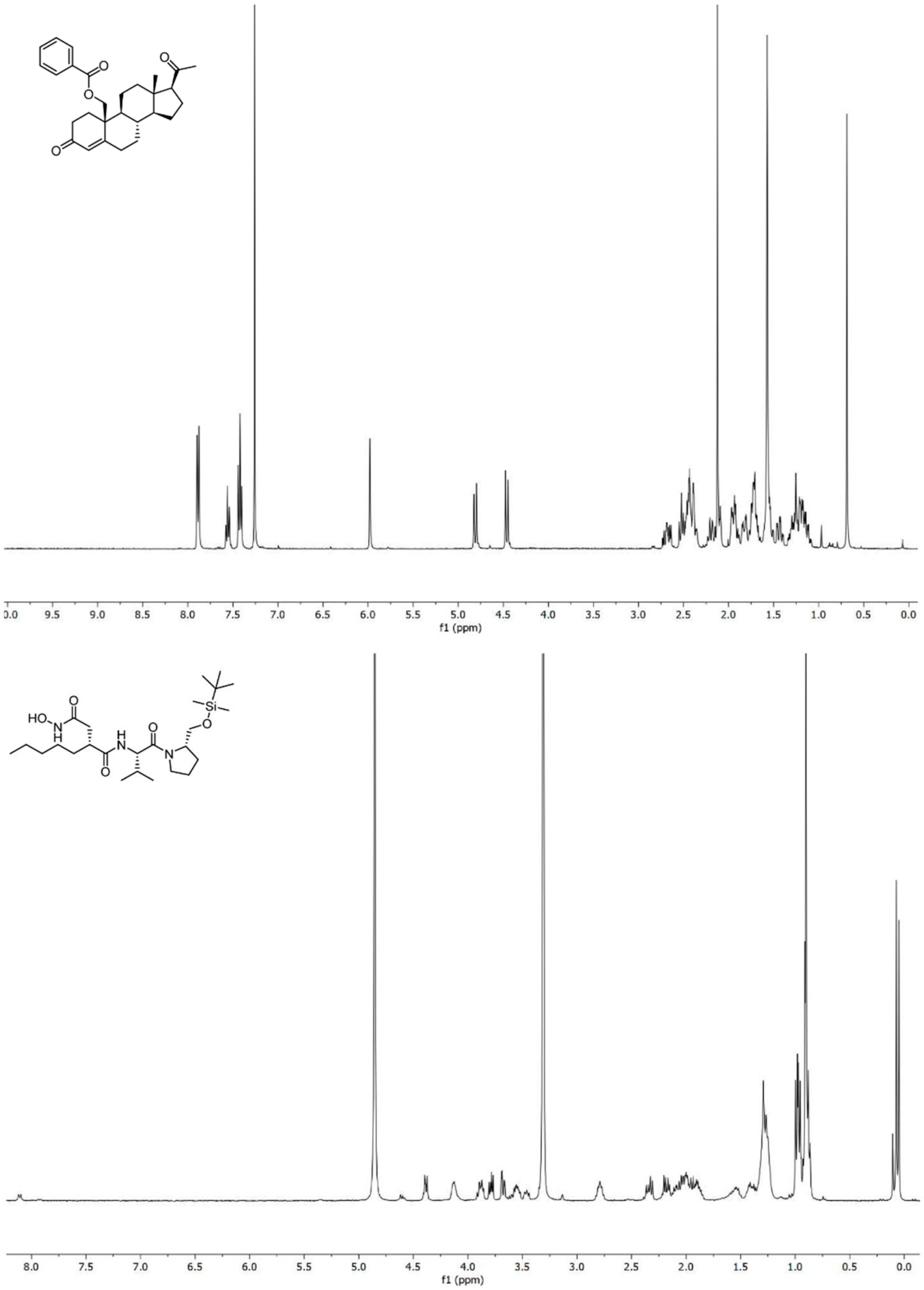


**Fig. S14: Chemical synthesis and $^1$H NMR spectra validation. A.** Chemical synthesis reaction of 19-O-Benzoyl-Progesterone (OBz-Progesterone, top) and Tertbutyldimethylsilyl-Actinonin (TBDMS-Actinonin, bottom). **B.** $^1$H NMR spectra of OBz-Progesterone (top) and TBDMS-Actinonin (bottom).

**Table S1: Metrics and cutoffs for binder design with MaSIF-seed.**

| Target | Motif | Site | Interface cutoff | NN score cutoff | Descriptor distance cutoff | #seeds | #selected seeds | #designs (#grafted seeds) | #select designs (#seeds) | Total design |
|---|---|---|---|---|---|---|---|---|---|---|
| Bcl2: Ven | S | 1 | 0.65 | 0.9 | 2.0 | 1743 | 78 | 28396 (69) | 1456 (67) | 1995 |
| | | 2 | 0.65 | 0.87 | 2.2 | 1048 | 33 | 7485 (29) | 464 (27) | |
| | | 3 | 0.6 | 0.85 | 2.3 | 1012 | 11 | 1073 (8) | 75 (8) | |
| DB3: Pro | H | 1 | 0.75 | 0.9 | 1.8 | 995 | 49 | 160488 (49) | 975 (44) | 1998 |
| | | 2 | 0.65 | 0.9 | 2.1 | 1046 | 36 | 147940 (36) | 548 (34) | |
| | S | 1 | 0.8 | 0.9 | 1.7 | 1775 | 98 | 10097 (39) | 475 (37) | |
| PDF1: Act | H | 1 | 0.65 | 0.87 | 2.2 | 1272 | 74 | 56813 (67) | 1447 (66) | 1997 |
| | S | 1 | 0.65 | 0.85 | 2.3 | 1373 | 98 | 3711 (56) | 550 (55) | |

**Table S2: Deep sequencing analysis of FACS-enriched populations.**

| Target | Design | Original scaffold | Scaffold set | Topology | Binding counts | Non-binding counts | Enrichment |
|---|---|---|---|---|---|---|---|
| Bcl2 | DBVen1619 | bGC_85 | Ref71 | EEHEE | 459449 | 15561 | 1.47 |
| DB3 | DBPro1156 | 3hC_242_0001 | Ref70 | HHH | 134116 | 652 | 2.31 |
| PDF1 | DBAct553 | 3hC_605_0001 | Ref70 | HHH | 35195 | 557 | 1.80 |

**Table S3: Computational analysis of ligand contributions.** Summary of different metrics in absence or presence of ligands. The buried solvent-accessible surface area (SASA) of the protein-ligand target complex, the percentage of ligand contribution to this buried SASA, computed binding energy (ddG) in Rosetta Energy Unit (R.E.U.) and the number of atoms in contact with the target complex have been measured. Atom contacts were calculated based on the Van der Waals radii ($r_{VdW}$ + 0.2Å tolerance) of each pair of atoms. N/A: Not applicable.

<table>
<tr><th>Design</th><th>Ligand</th><th>Target buried SASA [Å$^2$]</th><th>Ligand contribution</th><th>ddG [R.E.U]</th><th>ddG shift</th><th>Atom contact [Counts]</th><th>Atom cont. shift</th></tr>
<tr><td rowspan="2">DBVen1619_1</td><td>-</td><td>N/A</td><td>N/A</td><td>-25.94</td><td rowspan="2">-17.04%</td><td>94</td><td rowspan="2">+7</td></tr>
<tr><td>+</td><td>672</td><td>9.98%</td><td>-30.36</td><td>101</td></tr>
<tr><td rowspan="2">DBPro1156_1</td><td>-</td><td>N/A</td><td>N/A</td><td>-28.59</td><td rowspan="2">-18.46%</td><td>117</td><td rowspan="2">+7</td></tr>
<tr><td>+</td><td>702</td><td>10.57%</td><td>-33.87</td><td>124</td></tr>
<tr><td rowspan="2">DBAct553_1</td><td>-</td><td>N/A</td><td>N/A</td><td>-32.18</td><td rowspan="2">-27.72%</td><td>123</td><td rowspan="2">+26</td></tr>
<tr><td>+</td><td>886</td><td>12.13%</td><td>-41.10</td><td>149</td></tr>
</table>

**Table S5: Docking benchmark complexes.** List of 14 protein-ligand complexes and 200 decoys used in the binding partner recovery experiment. Search parameters: interface cutoff = 0.0; NN score cutoff = 0.8; descriptor distance cutoff = 3.0; #sites = 3; selection radius = 10Å.

| **Protein-protein-ligand complex** | | | |
|---|---|---|---|
| **PDB ID** | **Protein 1** | **Protein 2** | **Ligand** |
| 1A7X | FKBP12 (chain A) | FKBP12 (chain B) | BENZYL-CARBAMIC ACID [8-DEETHYL-ASCOMYCIN-8-YL]ETHYL ESTER (FKA) |
| 1S9D | ADP-Ribosylation Factor 1 (chain A) | Arno (chain E) | 1,6,7,8,9,11A,12,13,14,14A-DECAHYDRO-1,13-DIHYDROXY-6-METHYL-4H-CYCLOPENT[F]OXACYCLOTRIDECIN-4-ONE (AFB) |
| 1TCO | SERINE/THREONINE PHOSPHATASE B2 (chains A and B) | FK506-BINDING PROTEIN (chain C) | 8-DEETHYL-8-[BUT-3-ENYL]-ASCOMYCIN (FK5) |
| 3QEL | NMDA glutamate receptor subunit (chain A) | Glutamate [NMDA] receptor subunit epsilon-2 (chain B) | 4-[(1R,2S)-2-(4-benzylpiperidin-1-yl)-1-hydroxypropyl]phenol (QEL) |
| 4DRI | Peptidyl-prolyl cis-trans isomerase FKBP5 (chain A) | Serine/threonine-protein kinase mTOR (chain B) | RAPAMYCIN IMMUNOSUPPRESSANT DRUG (RAP) |
| 4MDK | Ubiquitin-conjugating enzyme E2 R1 (chain A) | Ubiquitin (chain E) | 4,5-dideoxy-5-(3',5'-dichlorobiphenyl-4-yl)-4-[(methoxyacetyl)amino]-L-arabinonic acid (U94) |
| 6ENG | DNA gyrase subunit B (chain A) | DNA gyrase subunit B (chain B) | Coumermycin A1 (BHW) |
| 6H0F | Protein cereblon (chain B) | DNA-binding protein Ikaros (chain C) | S-Pomalidomide (Y70) |
| 6N4N | NS3 protease (chain A) | Rosetta-designed danoprevir/NS3a complex reader 2 (chain F) | (2R,6S,12Z,13aS,14aR,16aS)-6-[(tert-butoxycarbonyl)amino]-14a-[(cyclopropylsulfonyl)carbamoyl]-5,16-dioxo-1,2,3,5,6,7,8,9,10,11,13a,14,14a,15,16,16a-hexadecahydrocyclopropa[e]pyrrolo[1,2-a][1,4]diazacyclopentadecin-2-yl 4-fluoro-2H-isoindole-2-carboxylate (TSV) |
| 6OB5 | Maltodextrin-binding protein (chain B) | Ankyrin Repeat Domain (AR), S3-2D variant | FARNESYL DIPHOSPHATE (FPP) |

| | | | |
|---|---|---|---|
| | | (chain D) | |
| 6QTL | VHH (chain A) | VHH (chain C) | CAFFEINE (CFF) |
| 6SJ7 | DDB1- and CUL4-associated factor 15 (chain A) | RNA binding protein 39 (chain C) | N~1~-(3-chloro-1H-indol-7-yl)benzene-1,4-disulfonamide (EF6) |
| 7DC8 | Switch Ab Fab light & heavy chain (chains A and B) | Interleukin-6 receptor subunit alpha (chains C and F) | ADENOSINE-5'-TRIPHOSPHATE (ATP) |
| 7TE8 | DB21 (chain A) | CA14 (chain C) | cannabidiol (P0T) |
| **Decoys** | | | |
| 1A2K_AB, 1A2K_C, 1AVX_A, 1AVX_B, 1BRS_A, 1BRS_D, 1ERN_A, 1ERN_B, 1H6K_B, 1H6K_Y, 1I07_A, 1I07_B, 1I4O_B, 1I4O_D, 1ID5_H, 1ID5_L, 1JKG_A, 1JKG_B, 1JZO_A, 1JZO_B, 1LQM_E, 1LQM_F, 1NPO_A, 1NPO_C, 1O9Y_A, 1O9Y_D, 1PXV_A, 1PXV_C, 1Q5H_A, 1Q5H_B, 1SHY_A, 1SHY_B, 1SOT_A, 1SOT_C, 1T0F_A, 1T0F_B, 1TQ9_A, 1TQ9_B, 1UGH_E, 1UGH_I, 1UUG_A, 1UUG_B, 1XDT_R, 1XDT_T, 1XPJ_A, 1XPJ_D, 1XT9_A, 1XT9_B, 1XUA_A, 1XUA_B, 1YC0_A, 1YC0_I, 1YLQ_A, 1YLQ_B, 1YY9_A, 1YY9_D, 1Z0K_A, 1Z0K_C, 1ZR0_A, 1ZR0_B, 1ZVN_A, 1ZVN_B, 2A2L_B, 2A2L_C, 2AQX_A, 2AQX_B, 2B3Z_C, 2B3Z_D, 2B42_A, 2B42_B, 2FE8_A, 2FE8_C, 2G2W_A, 2G2W_B, 2GD4_B, 2GD4_C, 2GKW_A, 2GKW_B, 2HDP_A, 2HDP_B, 2HEK_A, 2HEK_B, 2I32_A, 2I32_E, 2J12_A, 2J12_B, 2JI1_C, 2JI1_D, 2LBU_D, 2LBU_E, 2O8Q_A, 2O8Q_B, 2P45_A, 2P45_B, 2P47_A, 2P47_B, 2QLC_B, 2QLC_C, 2WAM_A, 2WAM_C, 2WQ4_A, 2WQ4_C, 2Y32_B, 2Y32_D, 2YZJ_A, 2YZJ_C, 2Z0P_C, 2Z0P_D, 2Z29_A, 2Z29_B, 2Z7F_E, 2Z7F_I, 3AXY_B, 3AXY_D, 3B5U_J, 3B5U_L, 3BTV_A, 3BTV_B, 3CDW_A, 3CDW_H, 3CEW_C, 3CEW_D, 3CG8_B, 3CG8_C, 3CHW_A, 3CHW_P, 3E2U_A, 3E2U_E, 3ECY_A, 3ECY_B, 3EYD_C, 3EYD_D, 3F74_A, 3F74_B, 3FJS_C, 3FJS_D, 3HCG_A, 3HCG_C, 3HN6_B, 3HN6_D, 3HRD_E, 3HRD_H, 3IBM_A, 3IBM_B, 3ISM_A, 3ISM_B, 3K3C_A, 3K3C_B, 3KMT_A, 3KMT_B, 3KZH_A, 3KZH_B, 3M85_B, 3M85_E, 3OGF_A, 3OGF_B, 3P71_C, 3P71_T, 3P8B_C, 3P8B_D, 3PGA_1, 3PGA_4, 3Q0Y_B, 3Q0Y_C, 3Q87_A, 3Q87_B, 3Q9U_A, 3Q9U_C, 3QWN_I, 3QWN_J, 3QWQ_A, 3QWQ_B, 3RDZ_A, 3RDZ_C, 3S8V_A, 3S8V_X, 3S9C_A, 3S9C_B, 3SGB_E, 3SGB_I, 3SLH_A, 3SLH_B, 3TND_B, 3TND_D, 3WN7_A, 3WN7_B, 4AG2_A, 4AG2_C, 4CJ0_A, 4CJ0_B, 4KGG_A, 4KGG_C, 4M5F_A, 4M5F_B, 4TQ1_A, 4TQ1_B, 4YDJ_G, 4YDJ_HL, 5GPG_A, 5GPG_B | | | |

**Table S5: Target protein and binder sequences.**

| Design | Sequence | Mutations from native |
|---|---|---|
| DBVen1619_1 | KYMLVVKGPNVTIFRWVDSSSEAETLARKIAKKLGLEVKSVEKKGNAVRVEIG | |
| DBVen1619_2 | QYLLVVKGPNVTKFRWVDSSSEAETLARKIAKKLGLEVKSVEKKGNAVRVEIG | K1Q, M3L, I13K |
| DBPro1156_1 | DEKAKTAETLIYQLFSKAMQQSDPNEAEKLLKKAEELAKKANDPRLEQVVRQHQVVV<br>RFLV | |
| DBPro1156_2 | DEKAKTAETLIWQLFGKAMQQSDPNEAEKLLKKAEELAKKANDPRLEQVVRQHQVVV<br>RFLV | Y12W, S16G |
| DBAct553_1 | DYIRELRAALILLALKKQHAEDPDAQRVADELMKKLFDAAHRNDKDKVKKVVEEAKKV<br>VSTY | |
| DBAct553_2 | DYIRELNRALILLALKKQHAEDPDAQRVADELMKKLFDAAHRNDKDKVKKVVEEAKKV<br>VSTY | R7N, A8R |
| DB3_H | QIQLVQSGPELKKPGETVKISCKASGYAFTNYGVNWVKEAPGKELKWMGWINIYTGE<br>PTYVDDFKGRFAFSLETSASTAYLEINNLKNEDTATYFCTRGDYVNWYFDVWGAGTT<br>VTVSSAKTTPPSVYPLAPGSAAQTNSMVTLGCLVKGYFPEPVTVTWNSGSLSSGVHT<br>FPAVLQSDLYTLSSSVTVPSSPRPSETVTCNVAHPASSTKVDKKIVPR | |
| DB3_L | DVVMTQIPLSLPVNLGDQASISCRSSQSLIHSNGNTYLHWYLQKPGQSPKLLMYKVSN<br>RFYGVPDRFSGSGSGTDFTLKISRVEAEDLGIYFCSQSSHVPPTFGGGTKLEIKRADA<br>APTVSIFPPSSEQLTSGGASVVCFLNNFYPKDINVKWKIDGSERQNGVLNSWTDQDS<br>KDSTYSMSSTLTLTKDEYERHNSYTCEATHKTSTSPIVKSFNR | |
| DB3_H<br>(Chimeric) | QIQLVQSGPELKKPGETVKISCKASGYAFTNYGVNWVKEAPGKELKWMGWINIYTGE<br>PTYVDDFKGRFAFSLETSASTAYLEINNLKNEDTATYFCTRGDYVNWYFDVWGAGTT<br>VTVSSASTKGPSVFPLAPSSKSTSGGTAALGCLVKDYFPEPVTVSWNSGALTSGVHT<br>FPAVLQSSGLYSLSSVVTVPSSSLGTQTYICNVNHKPSNTKVDKKVEPKSCDKTHT | |
| DB3_L<br>(Chimeric) | DVVMTQIPLSLPVSLGEQASISCRSSQSLIHSNGNTYLHWYLQKPGQSPKLLMYKVSN<br>RFYGVPDRFSGSGSGTDFTLKISRVEAEDLGIYFCSQSSHVPPTFGGGTKLEIKRTVA<br>APSVFIFPPSDEQLKSGTASVVCLLNNFYPREAKVQWKVDNALQSGNSQESVTEQDS<br>KDSTYSLSSTLTLSKADYEKHKVYACEVTHQGLSSPVTKSFNRGEC | |
| Anti-kappa_H | EVKLLESGGGLVQPGRSLRLSCIASGFDFSGYWMTWVRQAPGKGLEWIGDINPDSST<br>INSTPSLKDKVIISRDNAKNTLFLQMSKVRSEDTALYYCAQRGNYVPFPYWGQGTLVT<br>VSAAKTTPPSVYPLAPGSAAQTNSMVTLGCLVKGYFPEPVTVTWNSGSLSSGVHTFP<br>AVLQSDLYTLSSSVTVPSSTWPSETVTCNVAHPASSTKVDKKIVPRDCGCK | |
| Anti-kappa_L | SIVMTQTPKFLFVSAGDRVTITCKASQSVSNDVEWYQQKPGQSPKLMIYFASKRYNG<br>VPDRFTGSGFGTEFTFTISTVQAEDLAVYFCQQDYSSPWTFGGGTKLEIKRADAAPTV<br>SIFPPSSEQLTSGGASVVCFLNNFYPKDINVKWKIDGSERQNGVLNSWTDQDSKDST<br>YSMSSTLTLTKDEYERHNSYTCEATHKTSTSPIVKSFNRGEC | |
| Bcl2 | MAHAGRTGYDNREIVMKYIHYKLSQRGYEWDAGDDAEENRTEAPEGTESEVVHRAL<br>RDAGDDFERRYRRDFAEMSSQLHLTPDTARQRFETVVEELFRDGVNWGRIVAFFEF<br>GGVMCVESVNREMSPLVDNIAEWMTEYLNRHLHTWIQDNGGWDAFVELYGPSMR | |
| PDF1 | AILNILEFPDPRLRTIAKPVEVVDDAVRQLIDDMFETMYEAPGIGLAATQVNVHKRIVVM<br>DLSEDKSEPRVFINPEFEPLTEDMDQYQEGCLSVPGFYENVDRPQKVRIKALDRDGN<br>PFEEVAEGLLAVCIQHECDHLNGKLFVDYLSTLKRDRIRKKLEKQHRQQA | |

**Table 6: Crystallographic data collection and refinement statistics.**

| DBAct553_1:Actinonin:PDF1 (PDB: 8S1X) | |
|---|---|
| **Data collection** | |
| Space group | $P2_12_12_1$ |
| Cell dimensions | |
| *a*, *b*, *c* (Å) | 49.44, 75.01, 83.16 |
| a, b, g(°) | 90.0, 90.0, 90.0 |
| Wavelength (Å) | 0.87313 |
| Resolution (Å) | 55.7 - 1.88 (1.96 - 1.88) |
| Unique reflections | 24990 (1266) |
| *R*merge | 0.044 (1.125) |
| *I* / s*I* | 15.0 (1.3) |
| CC1/2 | 0.999 (0.426) |
| Completeness (%) | 96.9 (99.6) |
| Redundancy | 4.3 (4.4) |
| **Refinement** | |
| Resolution (Å) | 55.7 - 1.88 |
| No. reflections | 24982 (2801) |
| $R_{work}$ / $R_{free}$ | 0.1838/0.2030 |
| No. atoms | 1999 |

| | |
|---|---|
| Protein | 1850 |
| Ligand/ion | 73 |
| Water | 76 |
| *B*-factors (Å$^2$) | 57.2 |
| Protein | 56.9 |
| Ligand/ion | 66.7 |
| Water | 55.3 |
| R.m.s. deviations | |
| Bond lengths (Å) | 0.007 |
| Bond angles (°) | 0.730 |
| Ramachandran plot | |
| Favored (%) | 99.11 |
| Allowed (%) | 0.89 |
| Outliers (%) | 0.00 |
